# DeepTracer: Fast Cryo-EM Protein Structure Modeling and Special Studies on CoV-related Complexes

**DOI:** 10.1101/2020.07.21.214064

**Authors:** Jonas Pfab, Nhut Minh Phan, Dong Si

## Abstract

Information about macromolecular structure of protein complexes such as SARS-CoV-2, and related cellular and molecular mechanisms can assist the search for vaccines and drug development processes. To obtain such structural information, we present DeepTracer, a fully automatic deep learning-based method for fast de novo multi-chain protein complex structure determination from high-resolution cryo-electron microscopy (cryo-EM) density maps. We applied DeepTracer on a previously published set of 476 raw experimental density maps and compared the results with a current state of the art method. The residue coverage increased by over 30% using DeepTracer and the RMSD value improved from 1.29Å to 1.18Å. Additionally, we applied DeepTracer on a set of 62 coronavirus-related density maps, among them 10 with no deposited structure available in EMDataResource. We observed an average residue match of 84% with the deposited structures and an average RMSD of 0.93Å. Additional tests with related methods further exemplify DeepTracer’s competitive accuracy and efficiency of structure modeling. DeepTracer allows for exceptionally fast computations, making it possible to trace around 60,000 residues in 350 chains within only two hours. The web service is globally accessible at https://deeptracer.uw.edu.

## 1 Introduction

The determining factor for a protein’s functionality is its structure, which is given by a unique sequence of amino acids and its three-dimensional arrangement [1]. Consequently, researchers can draw conclusions about the behavior of the protein based solely on its molecular structure. This outcome can be useful in developing new vaccines and drugs as viral fusion proteins play a central role in how the viruses invade the host’s cells [2]. In order to prevent infections, researchers attempt to develop vaccines and medicines that target these fusion proteins. This strategy is currently applied to find an effective vaccine for the SARS-CoV-2 virus [3, 4, 5]. The structural information about the fusion proteins is crucial for researchers to predict their behaviors and ultimately to find the right vaccine [6].

To determine the structure of a protein, this work builds upon cryo-electron microscopy (cryo-EM) data [7]. Cryo-EM allows researchers to capture macromolecules’ three-dimensional maps, which describe the density of electrons at a near-atomic resolution. The technology has gained popularity in recent years as an alternative to established structure determination methods, such as X-ray crystallography, due to its improved quality and efficiency [8, 9]. Amid the current global crisis, it is important that cryo-EM is being deployed right alongside X-ray crystallography to support the search for medicines and vaccines to fight the current COVID-19 pandemic [10]. To derive the structure of a protein based on its 3D cryo-EM electron density map, researchers currently either have to manually fit the atoms or resort to existing template-based or homology modeling methods [11, 12, 13]. The manual fitting of atoms represents an enormous effort as proteins complexes usually consist of several thousand atoms, making it virtually impossible for larger structures. Therefore, there is a tremendous demand for a method that automatically determines the molecular structure from a cryo-EM density map. Unfortunately, existing tools [14, 15, 16, 17, 18] such as Rosetta, MAINMAST, and Phenix determine only fragments of a protein complex, or require extensive manual processing steps. Due to the ability of cryo-EM to capture multiple large proteins in the course of a single study [19, 20], a fully automated, efficient tool to determine complex structures would be crucial to increase the throughput of the technology and speed up the development of medicines.

In this paper, we present DeepTracer, a fully automated software tool that determines the all-atom structure of a protein complex based solely on its cryo-EM density map and amino acid sequence (Figure 1). No manual processing of the density map is necessary, and the tool requires no further parameters to run. The core of the method is a tailored deep convolutional neural network that allows for fast and accurate structure predictions when combined with complex pre- and post-processing steps. This paper significantly improved our previous preliminary method and results [21]. We also provide a web service and a CoV-related dataset along with the constructed models at DeepTracer’s website. To our knowledge, this is the first web service for fully-automated protein complex determination and coronavirus modeling using 3D cryo-EM.

**Figure 1:**
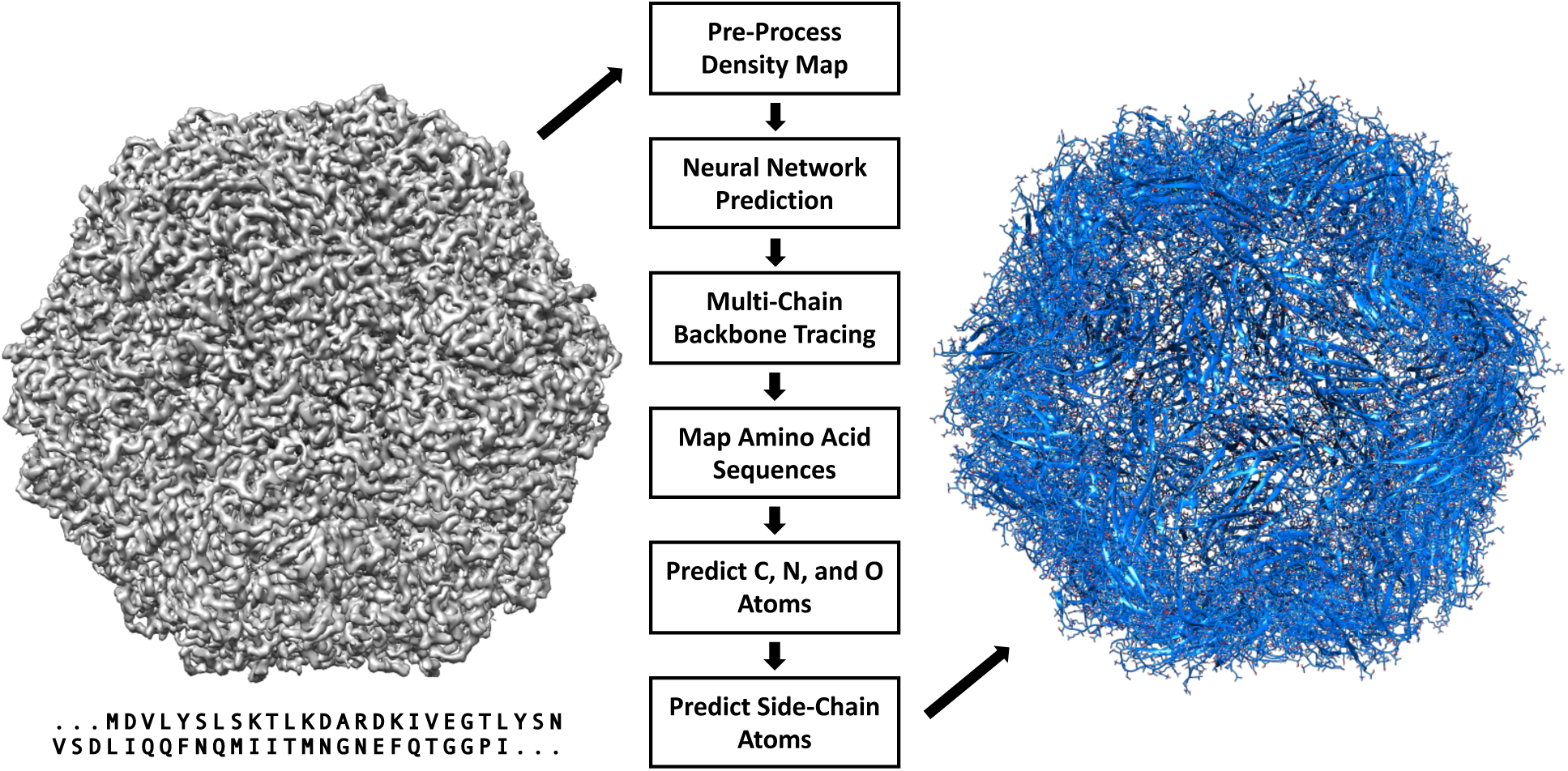
DeepTracer model determination pipeline. All-atom structure of multi-chain protein complexes is determined fully automatic solely from a density map and amino acid sequence using the steps shown in the center of the figure. The structure shown on the right side is an actual model built by DeepTracer.

## 2 Methods

DeepTracer performs an array of tasks to determine the structure of a protein. It pre-processes each density map for the neural network, feeds the density map to the network, and then transforms the output into a protein structure. An overview of the steps involved in this process is provided in Figure 1. In this section, we focus on all steps starting with the neural network. A detailed description of the pre-processing steps can be found in the supplementary material.

### 2.1 Neural Network Architecture

The convolutional neural network is the central piece of DeepTracer. Its job is to predict four vital pieces of information: the locations of amino acids, the location of the backbone, secondary structure positions, and amino acid types. Here, we take a closer look at the architecture of the neural network used in DeepTracer.

The U-Net forms the basis for DeepTracer’s neural network. It is a convolutional network architecture developed by researchers at the University of Freiburg. Its name derives from the U-shape of its architecture. The U-Net excels in fast and precise image segmentation tasks, particularly for biomedical applications [22]. For DeepTracer, we modified its original 2D architecture for 3D density maps and connected four separate U-Nets, one for each structural aspect (atoms, backbone, secondary structure elements, and amino acid types). The detailed architecture of the network used by DeepTracer can be seen in the in Figure 2. The pre-processed cryo-EM density maps are fed to the 64^3^ input layer of each U-Net. The output layer of each U-Net has the same 64^3^ shape with a varying number of channels depending on which structural aspect it predicts. The following paragraph describes the output channels in details.

**Figure 2:**
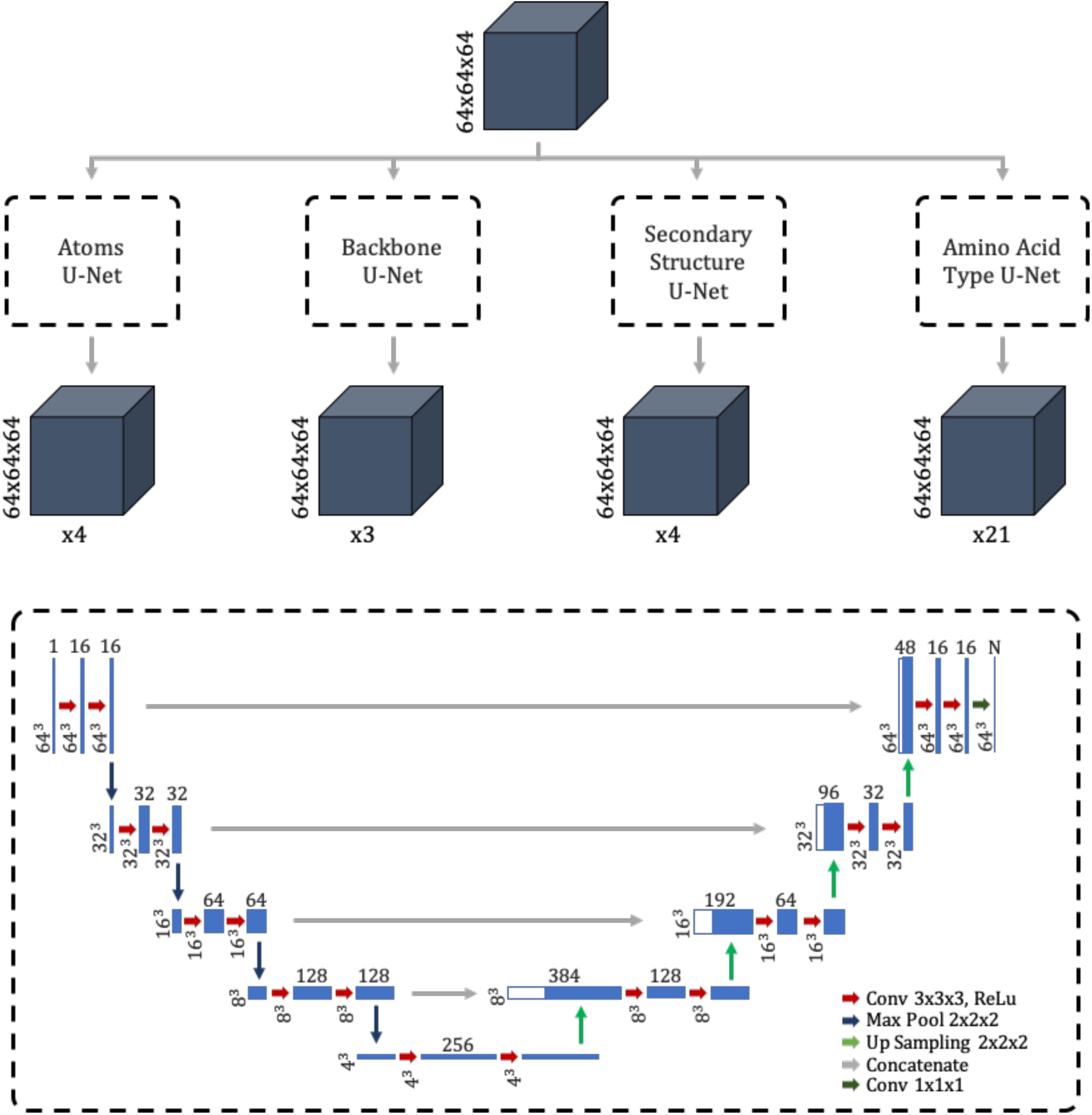
Architecture of tailored convolutional neural network. Top shows overview of DeepTracer’s neural network architecture consisting of four parallel U-Nets. The gray boxes show the input and output maps, with their dimensions noted to the left and the number of channels marked below. Bottom dashed box shows the detailed architecture of each parallel U-Net. The blue boxes show the output maps of the different layers where the dimensions of the maps are depicted on the left and the number of channels is depicted on top.

The overall convolutional network consists of four U-Nets. The first U-Net is the Atoms U-Net, which determines whether each voxel contains either a Cα atom, a nitrogen atom, a carbon atom, or no atom. Thus, its output has four channels. The second U-Net is the Backbone U-Net, which determines whether each voxel belongs to the backbone, meaning either it is on the backbone, a part of a side chain, or not a part of the protein. Thus, it has three different output channels. Next, the Secondary Structure U-Net is responsible for finding out the secondary structure of each voxel. It has four output channels for loops, sheets, helices, and no structure. Finally, the Amino Acid Type U-Net determines the amino acid type for each voxel. As 20 different types of amino acids have been found in nature, this U-Net has 21 output channels, representing the amino acids plus the case in which the voxel is not part of the protein.

### 2.2 Training Data Collection

Before training the U-Net model, we have to collect a training dataset. Previous projects, such as [17], used simulated density maps to train their neural networks. However, for the network to learn common noise patterns in cryo-EM density maps, we decided to use experimental maps. The maps were downloaded from the EM-DataResource website [23] together with their deposited model structures that served as the ground truth in the training process and were fetched from RCSB Protein Data Bank [24]. As this work focuses on high resolution maps, we only used density maps with a resolution of 4Å or better. In total, we downloaded 1,800 experimental density maps and their corresponding deposited model structures. The maps were randomly split into training and validation sets with an 80:20 ratio.

To label each density map, we created masks with the same dimensions as the grid of the density map, providing a label for each voxel. The labels of the masks were hereby created based on the deposited model structures of each density map. As shown in Figure 2, the model has four different outputs, for each of which we created separate masks. The atoms mask provides a label for each voxel whether or not it contains a Cα, C, or N atom. Therefore, we filtered out these atoms from the protein structure, calculated the corresponding grid indices for their location, and set that voxel and all directly neighboring voxels to the value representing the atom (1 for Cα, 2 for C and 3 for N atoms). A visualization of an atom mask can be found in the supplementary material in Figure S5.

The masks for the backbone, secondary structure, and Amino Acid Type U-Net, were created in a similar manner. The backbone mask filters all backbone atoms and side-chain atoms and sets the respective voxels and all surrounding voxels with a distance of 2 to 1 for backbone and 2 for side chain. To create the secondary structure mask, we filtered all atoms for helices, sheets, and loops and then set all voxels with a distance of 4 surrounding the atoms to 1 for loop, 2 for helix, and 3 for sheet. Finally, for the amino acid type mask, all Cα atoms for each of the 20 amino acid types were filtered out, and all surrounding voxels within a distance of 3 were set to a value between 1 and 20, where each value corresponds to a specific amino acid type. An example of all masks can be seen in Figure 3. An example of a raw prediction from the trained neural network for the EMD-6272 density map can be found in Figure S4 from the supplementary materials.

**Figure 3:**
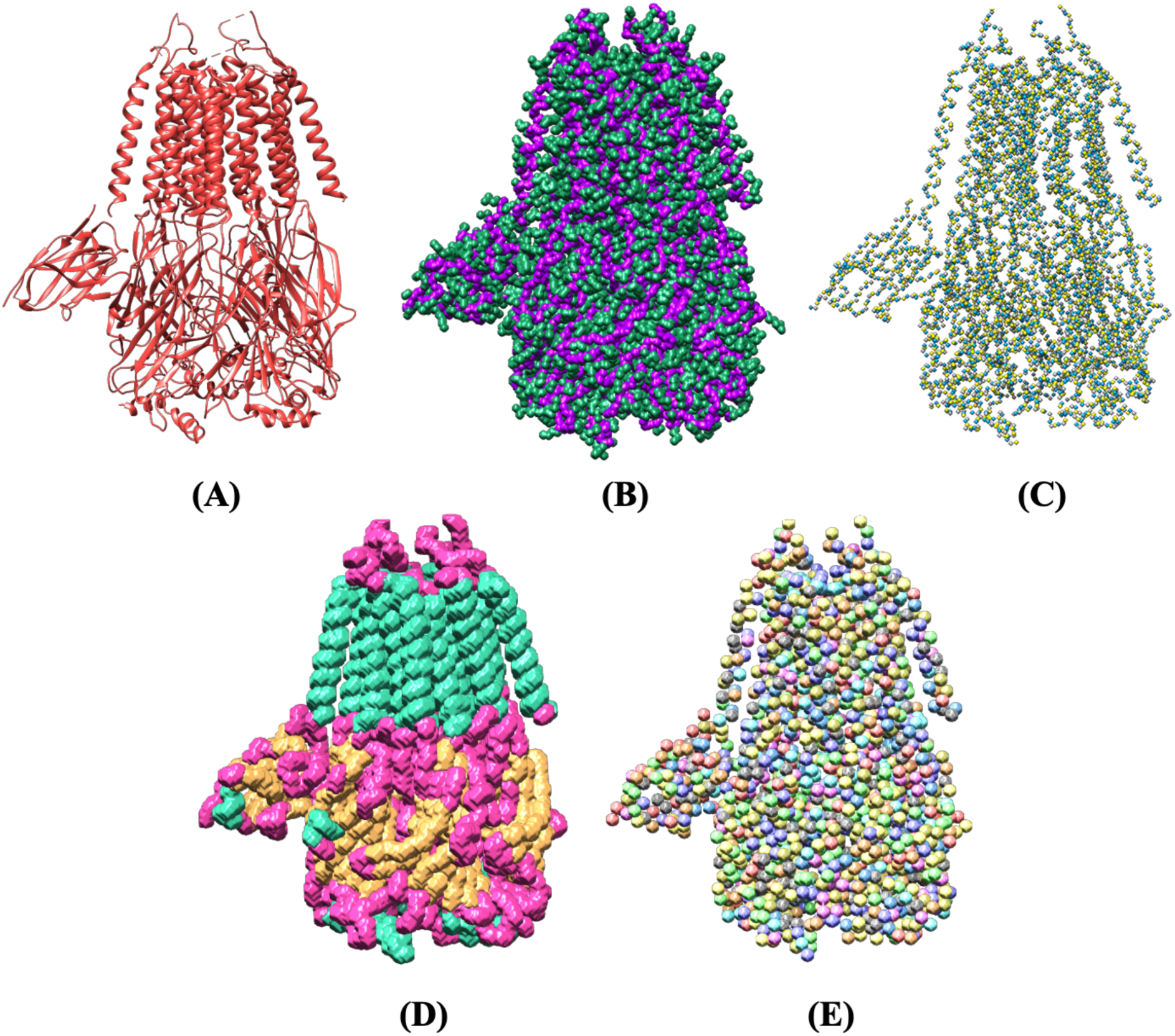
Example masks from the training dataset based on the PDB-6NQ1 deposited model structure. (A) Deposited model structure. (B) Backbone (Cα, C, and N atoms) in purple and side chains in green. (C) Atoms mask with labels for Cα, C, and N atoms. (D) Secondary structure mask with helices in turquoise, loops in pink and sheets in orange. (E) Amino acid type mask with 20 different colors.

### 2.3 Tracing Backbone

This step uses the output of the U-Net to create an initial model structure that contains only Cα atoms connected into chains. This is a central post-processing step, and its accuracy determines to a great extent how well the remaining post-processing steps will perform. The step can be split into three different parts: identifying disconnected chains, which can be processed independently; calculating the x,y, and z coordinates of the Cα atoms; connecting the Cα atoms into chains by applying a modified travelling salesman algorithm.

Identifying chains prior to any atom prediction has two advantages. First, it improves the performance of the step as each chain will contain a lower number of atoms that have to be connected by the travelling salesman algorithm. Second, it decreases the number of incorrect connections between atoms of separate chains as they are processed independently. To identify chains, we used the output of the Backbone U-Net. We rounded each voxel of the confidence map to either zero or one and then found connected areas of voxels with a value of one. Disconnected areas were then identified as separate chains. An example of the chain identification process visualized for the EMD-0478 density map can be seen in Figure 4.

**Figure 4:**
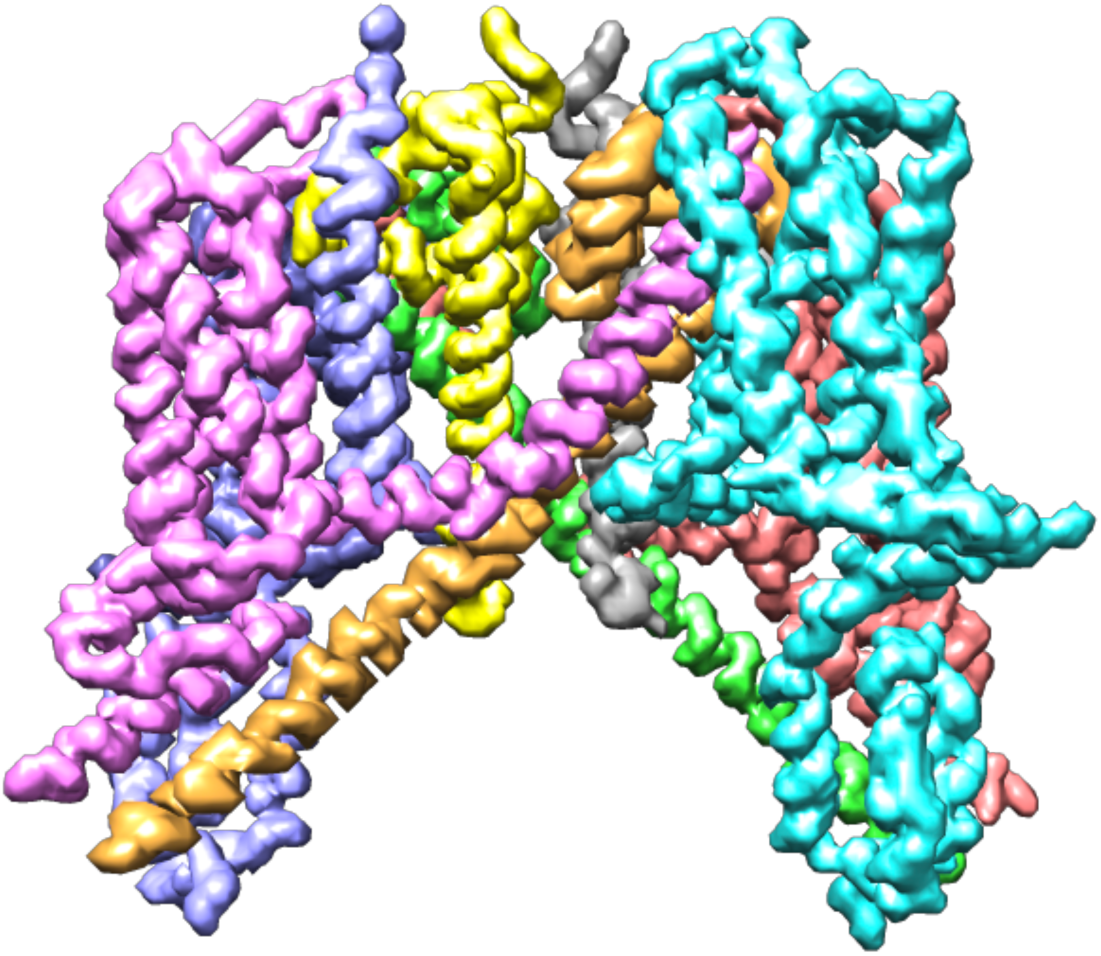
Backbone confidence map of the EMD-0478 density map with identified chains annotated in different colors.

To find the x, y, and z coordinates of the Cα atoms, we utilized the Cα channel from the output of the Atoms U-Net. A voxel value in this map describes the confidence of whether this voxel contains a Cα atom. The coordinates were then calculated in two steps. First, we found the indices of all local maximums in the confidence map within a distance of 4 voxels that have a minimum value of 0.5. Next, we refined the indices by calculating the center of mass of all voxels within a distance of 4 surrounding the local maximums. This is possible as we moved away from integer indices towards floating point coordinates, giving us the opportunity to express locations more precisely.

The most challenging part of this step is to connect the placed Cα atoms into chains correctly. The factorial growth of the number of ways in which the atoms can be connected makes it infeasible to test all possible solutions even for a low number of atoms. Therefore, we decided to solve the problem using an optimization algorithm, particularly, for the travelling salesman problem (TSP). However, our problem does not match every criterion of the traveling salesman problem. The shortest possible path is not necessarily the correct one as the ideal distance between Cα atoms is 3.8Å [25]. Deviations from this value are, however, possible due to prediction inaccuracies. Additionally, it is often difficult to decide only based on the distance which atoms to connect if there are multiple possibilities with a similar distance. To address these issues, we developed a custom confidence function instead of solely relying on the euclidean distance between atoms. The confidence function’s idea is to return a score between 0 and 1, which expresses how confident we are that these two atoms are connected. The goal of the TSP algorithm is then to connect the atoms such that the sum of all confidence scores between connected atoms is maximized.

The calculation of the confidence score between Cα atoms considers two factors: the Euclidean distance between the atoms, and the average density values of voxels that lay in between the atoms on the backbone confidence map predicted by the Backbone U-Net. The latter factor is to ensure that connections are made along the backbone of the structure. The voxels that lay between the atoms are found using Bresenham’s algorithm [26]. To transform these metric values to a confidence score, we used a probability density function *p*(*x, µ, σ*) with a mean μ, which represents the ideal metric value, and a standard deviation s. To make sure that the function returns exactly 1 at the mean, we normalized it by dividing it by the probability density value at the mean. For the euclidean distance, we used a mean of 3.8 and a standard deviation of 1. The average backbone confidence has a mean of 1 and a standard deviation of 0.3. The standard deviations were determined based on several rounds of testing. Both probability density functions can be seen in Figure S6. In order to combine both results into a single confidence score, we simply multiply both values. As the TSP algorithm was designed to minimize distances between paths, we then just subtract the confidence score from 1 and provide it to the algorithm.

To apply the TSP algorithm, we had to specify a start/end point. However, we could not know yet at which atom the chain will start and end. Therefore, we added a new atom that is connected to every other atom with a confidence of 1. This atom was then specified as the start/end and later on removed from the actual chain. An example of the application of the TSP on a list of Cα atoms can be seen in Figure 5.

**Figure 5:**
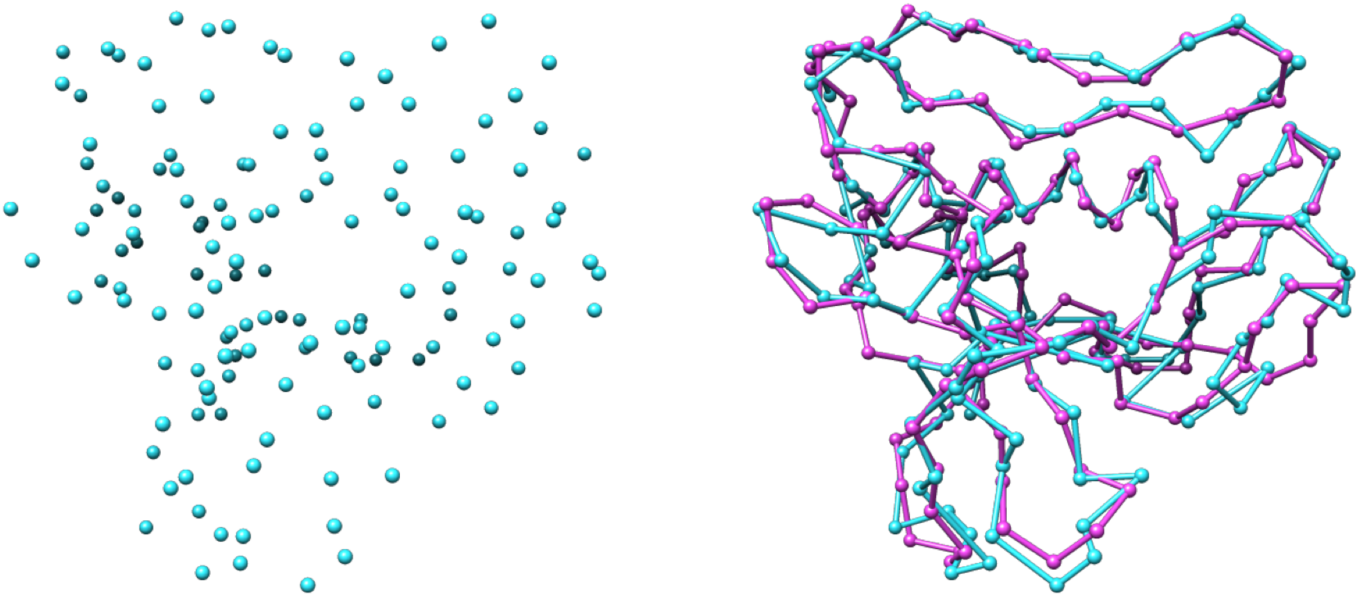
Traced backbone atoms. Predicted Cα atoms for the EMD-4054 density map in blue before (left) and after (right) the backbone tracing step compared to the deposited model structure in pink.

### 2.4 Amino Acid Sequence Mapping

To realize the side-chain prediction for the protein structure, we first need to know each amino acid’s type. As discussed in Section 2.1, one output of the deep learning model is the amino acid type prediction. However, depending on the resolution of the density map, this prediction is of limited accuracy with around 10% to 50% since some amino acids have a similar appearance in electron density maps. The goal of this step is to improve the amino acid type accuracy by aligning intervals of the initially predicted sequence to the known true amino acid sequence (protein primary structure) and then updating the types of the predicted amino acids accordingly (see Figure 6).

**Figure 6:**
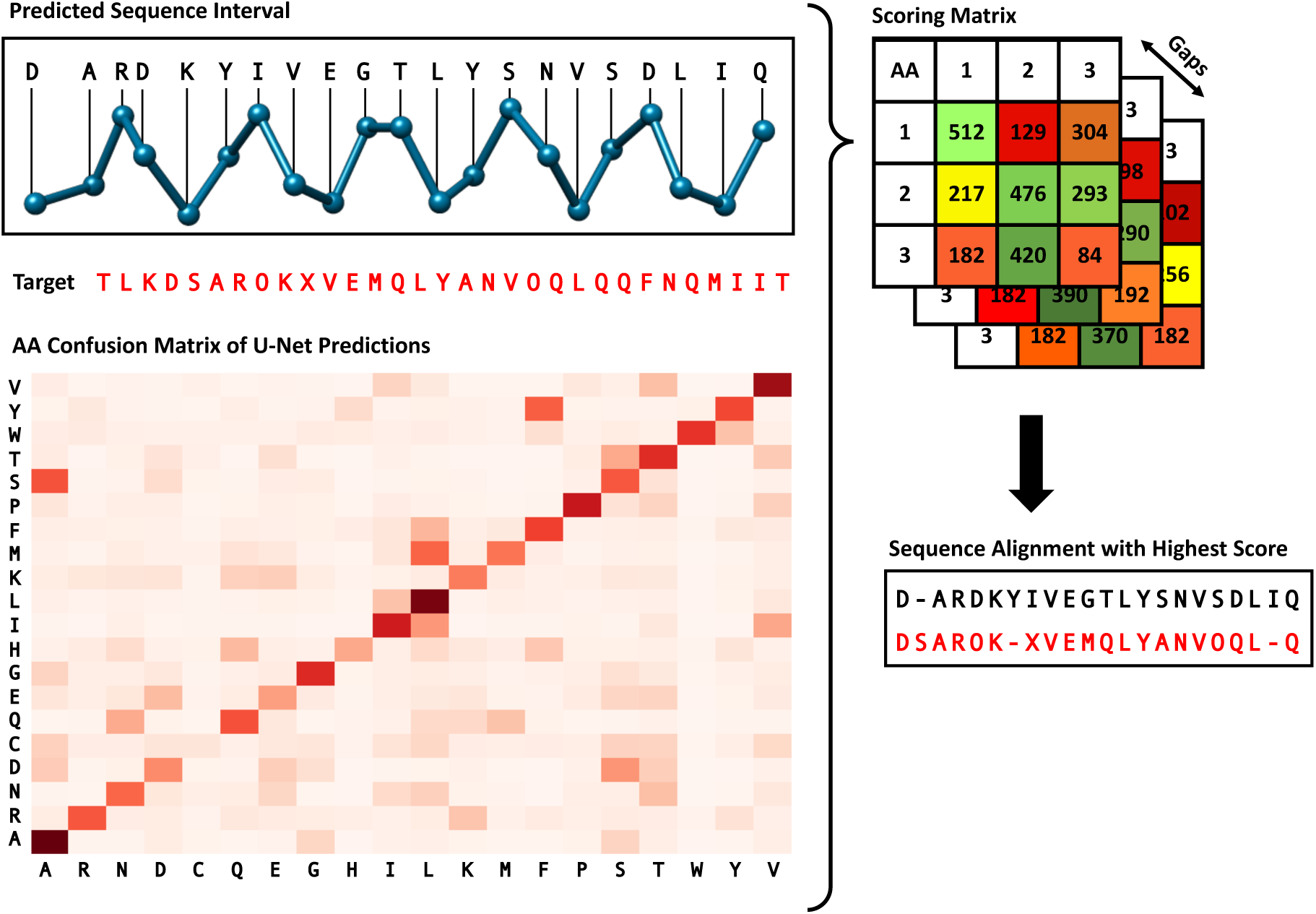
Protein sequence alignment algorithm. Interval of the predicted sequence is aligned with the target sequence using a custom dynamic algorithm. The amino acid confusion matrix depicts the relative frequency of pairs of predicted and true amino acid type and was calculated based on a set of test density maps. The numbers shown in the score matrix are solely for illustrative purposes and do not reflect real data.

Aligning amino acid sequences is a common problem in the field of bioinformatics, and previous research has led to the development of multiple algorithms [27, 28, 29]. However, these algorithms are usually applied between different proteins to measure their sequence similarities, which does not quite fit our use-case. The main problem is that we require an algorithm that does not treat all matches and mismatches in the same way. This stems from the fact that some amino acid types have a more similar appearance in density maps than others, which leads to some mismatches of the U-Net being more likely than others. To analyze the relative frequency of a certain match of predicted and true amino acid type, we applied the U-Net to 200 different density maps and compared the predicted amino acid types with the actual types from the deposited model structures. The heatmap depicting this analysis is shown in Figure 6. As expected, the most frequent matches are those of the same predicted and true amino acid type. However, we can also see that the U-Net often mixes up some types (e.g., ALA and SER) and struggles more with other types (e.g., CYS).

To incorporate the U-Net prediction behavior described in the previous section into the alignment algorithm, we defined a reward function *r* which returns a score denoting how valuable a certain match of predicted type *p* and true type *t* is. With *f* (*p, t*) defined as the relative frequency of a match, we constructed the reward function shown in Equation (1). The constant 100 as a multiplier is used to balance the match rewards with gap penalties described in the next section, and was chosen based on multiple rounds of testing. The 0.05 constant was chosen as this represents the likelihood of a correct match if we would chose the amino acid type randomly, since there are 20 different types of amino acids. The score is zero if the relative frequency equals this random likelihood.

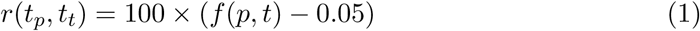

In addition to the match reward, our algorithm also requires a gap penalty. A gap represents a skipped amino acid in either the predicted or true sequence. This penalty, however, cannot simply be a static value as not all gaps are the same. For example, gaps in the beginning of a sequence before any matches were made should not result in any penalties as we only match short intervals of the predicted sequence, meaning it is highly unlikely that they align at the first amino acid of the true sequence. Additionally, the number of consecutive gaps is important. Cases where DeepTracer misses an amino acid or predicts an extra amino acid appear relatively frequent meaning that a single gap is not unlikely. However, two missed amino acids in a row is very uncommon, and three gaps in a row virtually never happens. Therefore, we must define our penalty function *p* such that it takes the number of consecutive gaps *g* into account. Let *i* be the index of the amino acid that is not skipped. Then we can define *p* as shown in Equation (2). The constants 20 and 30 were chosen based on test runs to create a good balance with the rewards function.

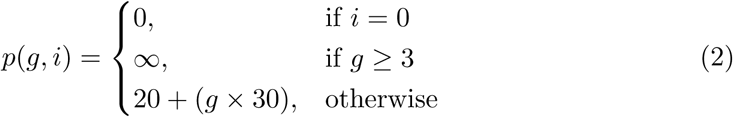

Since we have defined a reward and penalty function, we can find the ideal alignment by maximizing the sum of all rewards and penalties using a dynamic algorithm. To do so, we defined a recursive equation which calculates the optimal solution based on an index *i*, which points to the current amino acid in the true sequence, an index *j*, which points to the current amino acid in the predicted sequence, and *g*, which counts the number of previous consecutive gaps. With *t* and *p* as the true and predicted sequence, we defined this function as shown in Equation (3). To efficiently find the solution, we applied the dynamic programming “bottom up” approach [30].

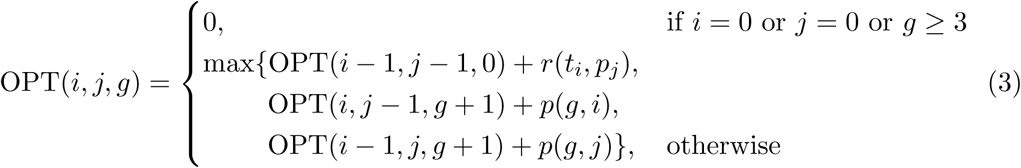

### 2.5 Carbon, Nitrogen, and Oxygen Determination

So far, the determined residues consist solely of Cα atoms. A complete protein backbone also consists of carbon, nitrogen, and oxygen atoms. Previous research has introduced various methods for reconstruction of a protein backbone from a reduced representation, such as one contains only Cα atoms [31]. Instead of employing these theoretical methods, we chose to implement our own backbone reconstruction method to make use of the information captured from the 3D cryo-EM density maps. This section presents our all-atom backbone reconstruction method. This is necessary for the next step in the pipeline, resolving the side-chain atoms.

In addition to Cα prediction, the U-Net also provides information about carbon and nitrogen atoms in the confidence map predicted by the U-Net. We can use this information in combination with the previously determined Cα atom positions to place the carbon and nitrogen atoms. Between the Cα atoms of two connected amino acids, there is always a nitrogen and carbon atom. Therefore, we can guess the initial position of these atoms by calculating the vector from one Cα atom to the other and then placing the nitrogen and carbon atoms at one third and two third of the distance of this vector. To refine these initial positions we calculated the center of mass around them in the carbon and nitrogen confidence maps. In Figure 7a we can see an example for the initial and refined placement of the carbon and nitrogen atoms.

**Figure 7:**
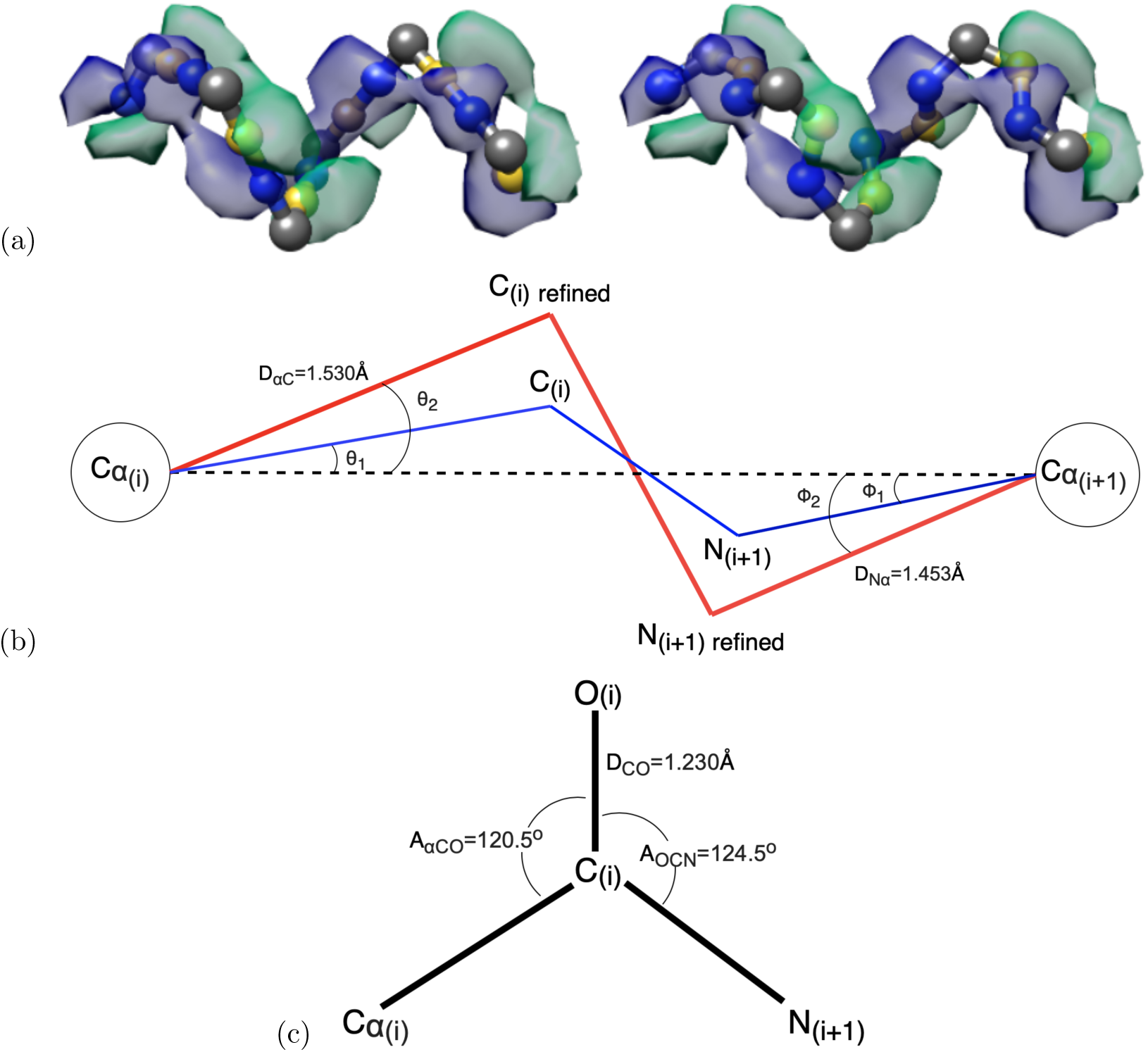
Carbon, nitrogen, and oxygen determination. **(a)**, Initial positioning of carbon (yellow) and nitrogen (blue) atoms in between the Cα atoms (gray) on the left and their initial refined positioning, which fits the U-Net prediction of carbon atoms (green volume) and nitrogen atoms (blue volume), on the right. **(b)**, The positions of carbon and nitrogen atoms are refined further by forcing bond angles into their well-known values. The blue lines represent the bonds from the initial refinement. The red lines represent the bonds from the final refinement. **(c)**, Position of oxygen atom in the carbonyl group by definition.

After the initial refinement, we can further refine the positions of the carbon and nitrogen atoms by applying well-known molecular mechanics of a peptide chain. We made several assumptions about the positions of carbon, nitrogen, oxygen atoms relative to the Cα atoms as seen in Figure 7b. First, we assumed the planar peptide geometry in which the Cα atom and carbon atom in the carbonyl group of an amino acid are in the same plane as the next amino acid’s nitrogen and Cα atom [32]. Second, we constructed a virtual bond between the neighboring Cα atoms. The angles between this bond and Cα_(*i*)_–C_(*i*)_ bond (ϑ_2_) and between this bond and Cα_(*i*+1)_–N_(*i*+1)_ bond (φ_2_) are 20.9° and 14.9°, respectively [32]. Third, the peptide bonds in a protein are in the stable trans configuration [33].

To refine the position of the carbon atoms, we relied on the previous refinement. Let us call the unit vector pointing from Cα_(*i*)_ to C_(*i*)*refined*_ *v*_1_, the unit vector pointing from Cα_(*i*)_ to C_(*i*)_ *v*_2_, and the unit vector pointing from Cα_(*i*)_ to Cα_(*i*+1)_ *v*_3_.

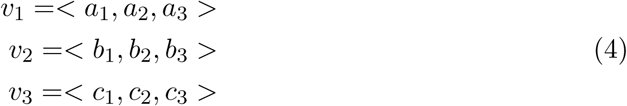

The goal is to solve for the components of v_1_. Due to the planar peptide geometry, *v*_1_, *v*_2_, and *v*_3_ exist in the same plane. Thus, their triple product equals to zero.

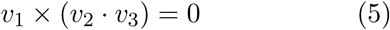

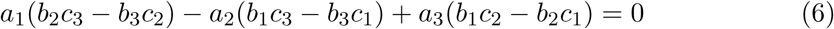

From this relation and the cross product of *v*_1_ and *v*_2_, and that of *v*_2_, *v*_3_, we can construct a system of equations:

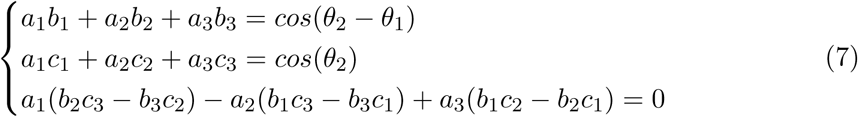

Solving this system of equation yields *a*_1_, *a*_2_ and *a*_3_. Next, the vector *v*_1_ is scaled appropriately to resolve the new position of the carbon atom. The position of the nitrogen atom is refined in a similar manner.

To determine the location of the oxygen atom in the carbonyl group, we assumed the coplanar relationship between the oxygen, Cα, carbon, and nitrogen atom [32], and that the angle A_*αCO*_ and A_*OCN*_ (see Figure 7c) are approximately identical. We then derived a unit vector pointing in the direction of the C-O bond and scale it with the C-O bond length to get the position of the oxygen atom.

### 2.6 Side Chain Prediction

The final step of DeepTracer is the side chain prediction. Its goal is to position the side chain atoms of each amino acid based on its type and backbone structure. This is done by using SCWRL4 [34], a tool developed by the Dunbrack lab, which predicts side chain atoms for structures that have a complete backbone and amino acid types set. The tool is integrated in the pipeline of DeepTracer and runs fully automatic as well. It also performs a collision detection to ensure that side-chains of different residues do not overlap. In Figure S7 we can see an example of an α-helix after the side chain prediction step.

## 3 Results

We evaluated the effectiveness of DeepTracer by applying it to multiple test datasets of experimental density maps, most of which depict multi-chain complexes. As a point of comparison, we used results generated by Phenix’s map-to-model function. Further comparative tests with Rosetta and MAINMAST method can be found in the supplementary materials.

### 3.1 Metrics

To ensure the objectivity of the comparison with the existing Phenix method, we used the `phenix.chain_comparison` tool [35], which is available at no cost as part of the Phenix software suite. This tool compares two models by finding a one-to-one matching between their residues based on Cα positions. For a two residues to match, they cannot be further apart from the other than 3Å. Based on this matching, several metrics are calculated. The first metric is the root-mean-square deviation (RMSD), which expresses the average distance between Cα atoms of matched residues. Second, the coverage is expressed using the matching percentage. This value represents the proportion of residues from the deposited model, which have a matching interpreted residue and is calculated by dividing the number of matches by the total number of residues. Third, to evaluate how well the amino acid types were predicted, the `chain_comparison` tool calculates the sequence matching percentage, which denotes the percentage of matched residues that have the same amino acid type. Lastly, to get a sense of how similar residues are connected, the mean length of matched segments is calculated where consecutive matches are connected in both models. Besides the metrics calculated by the `phenix.chain_comparison` tool, we also apply the LGA (Local-Global Alignment) algorithm, which aligns two models and computes the GDC (Global Distance Calculation) score. This score measures the similarity of two structures based on all atoms (including side-chains) on a range of 0 to 100 with 100 being a perfect match [36, 37]. We applied it on the most important dataset of SARS-CoV-2 density maps due to the high manual and computational effort involved in the calculation of this metric.

### 3.2 Phenix Benchmark Test

We applied DeepTracer on a set of 476 density maps assembled by the authors of Phenix’s map-to-model method [14] and compared the determined models against the ones published on Phenix’s website [38] (see Figure 8). We can see that DeepTracer achieves better results than the Phenix method for every metric calculated by the `phenix.chain_comparison` tool. The matching percentage of deposited model residues is, on average, 76.93% compared to 45.65% with Phenix, representing an improvement of over 30%. DeepTracer achieved a matching percentage above 70% for almost all density maps, except a few outliers. The average RMSD value of DeepTracer (1.29) is 0.11 higher than Phenix’s (1.18). We can see that the distribution of the RMSD values of DeepTracer follows a similar pattern as Phenix, with a strong correlation between RMSD and the resolution of the density map. The most significant improvements of DeepTracer were measured for the sequence matching which expresses the percentage of matched residues in the determined and deposited model that have the same amino acid type. For this metric, DeepTracer achieved 49.83%, which is more than four times higher than the 12.29% of the Phenix method. Although 49.83% is still fairly low, the distribution of the values shows that there is a steep improvement of the sequence matching with more accurate maps. Two factors contribute to this trend. First, side chain atoms, which determine the amino acid type, are only visible in very high resolution maps, making it almost impossible to accurately predict the amino acid type for lower resolutions. Second, the amino acid type mapping of every segment can either be correct or incorrect. It means that either all amino acid types will be correct for this segment, or in case of an incorrect mapping, the amino acid types are entirely random. This amplifies the steep incline in accuracy for higher resolution maps. Third, for the last evaluated metric, the mean length of matched segments, improved from 8.16 with Phenix to 14.05 with DeepTracer. While this number is influenced by several factors including the average length of connected segments in the deposited model structure, this is an indicator that DeepTracer connects residues better than the Phenix method.

**Figure 8:**
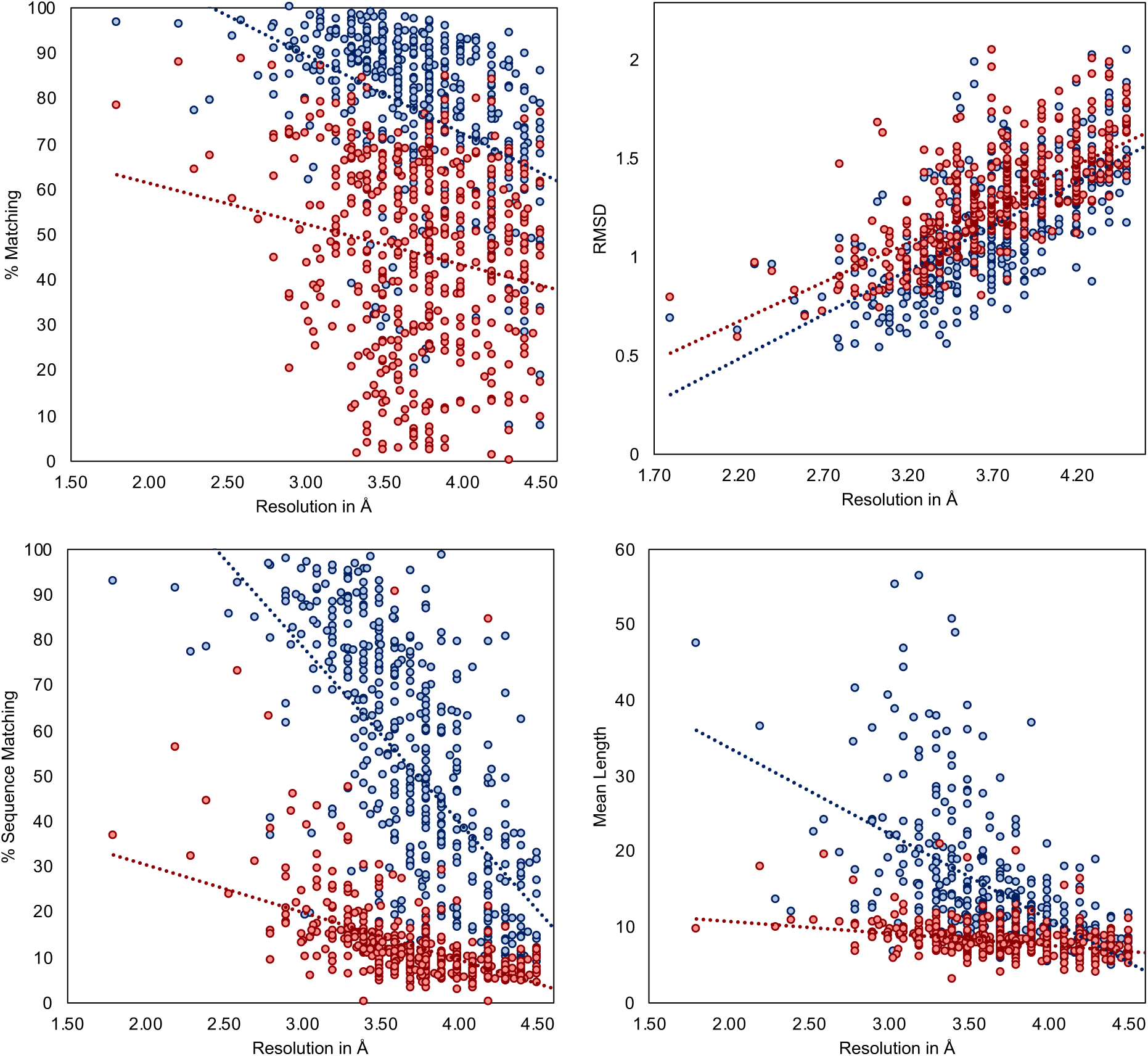
Evaluation results for set of 476 experimental density maps. Evaluation of determined models from DeepTracer (blue) and Phenix (red) for 476 density maps. The dotted lines represent the trend for each method. DeepTracer outperformed Phenix in all four metrics. Precise data can be found in Table S3.

Figures 9 and 10 show multi-chain complexes modelled by DeepTracer and Phenix compared to the deposited model structure. In both figures we can note that Deep-Tracer’s model is more complete. In Figure 9, we can particularly see the greater coverage and more precise placement of the residues in direct comparison with the ground truth. Figure 10 shows well that, even though Phenix determined most of the residues correctly, DeepTracer connected the residues better creating a less fragmented model.

**Figure 9:**
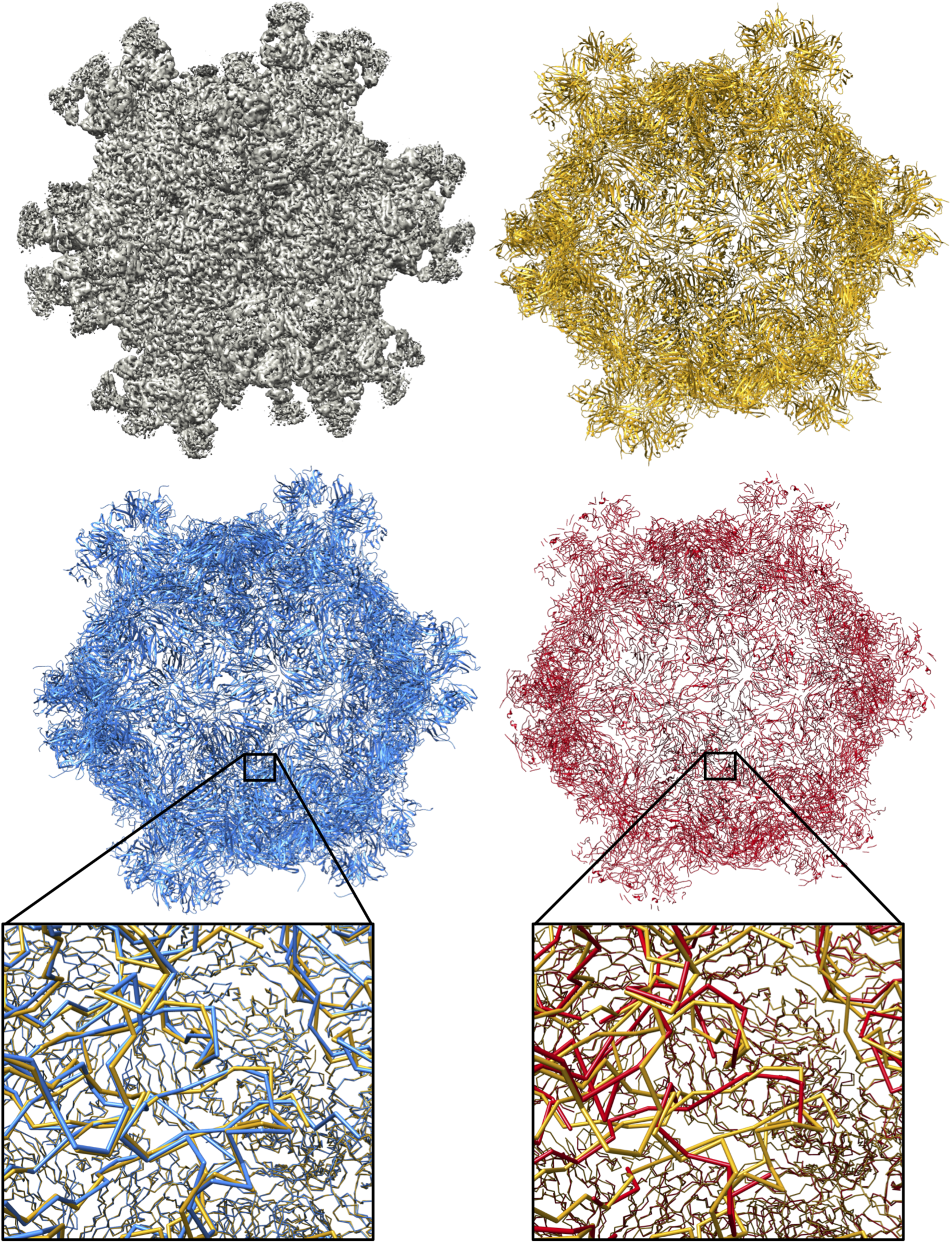
Results of EMD-6572 density map. Models built by DeepTracer (blue) and Phenix (red) next to PDB-3J9S deposited model structure (yellow) for EMD-6572 density map.

**Figure 10:**
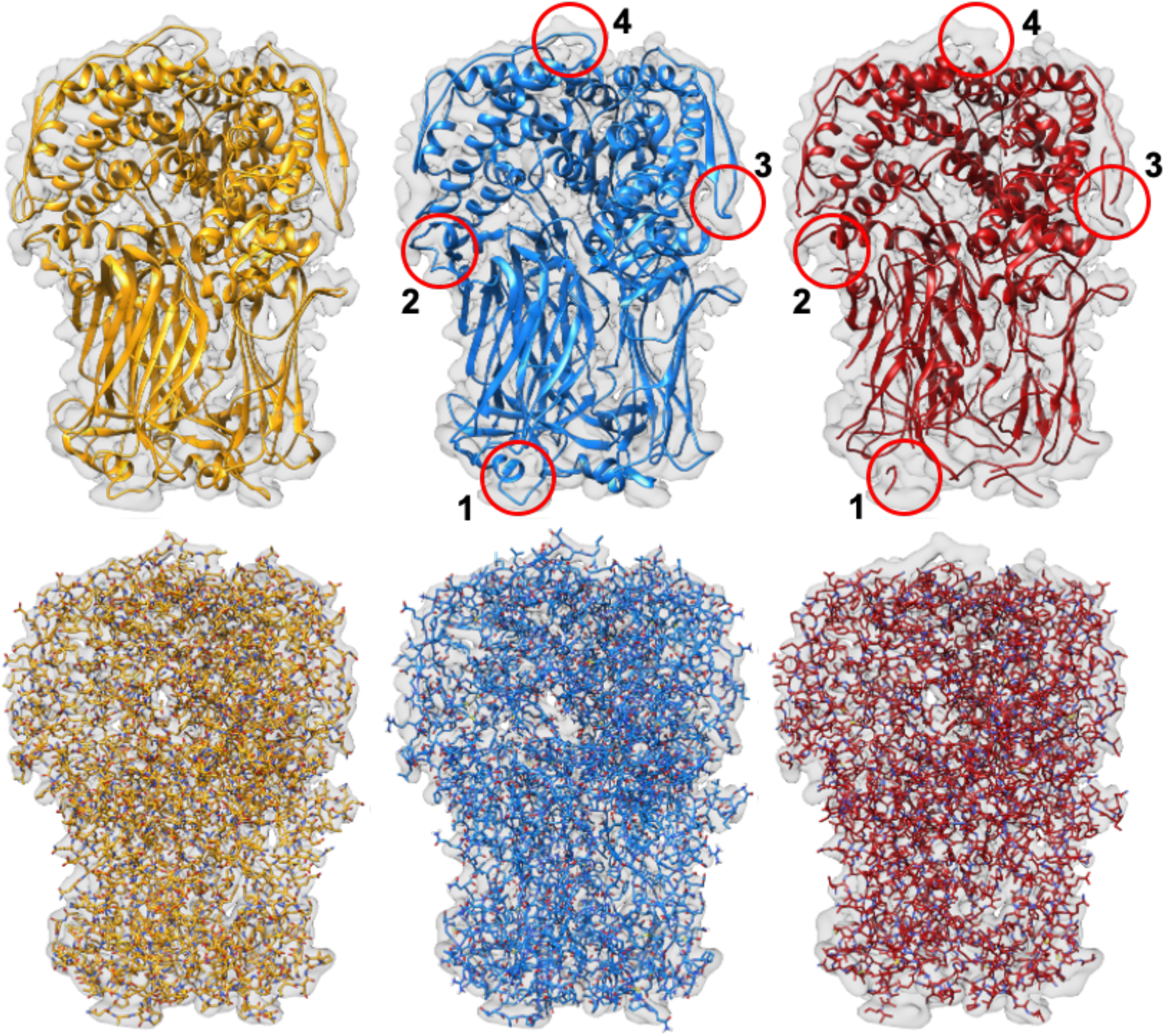
Results of EMD-6272 density map. Models built by DeepTracer (blue) and Phenix (red) compared to PDB-3J9S deposited model structure (yellow) for EMD-6272 density map. Top row shows structures in ribbon view and lower row in all-atom view.

### 3.3 Coronavirus-Related Results

In the search for an effective COVID-19 vaccine and medicine, structural information about the viral protein is crucial. Therefore, we applied DeepTracer on a set of coronavirus-related density maps to demonstrate how it can aid researchers in obtaining such structural information. To create a point of comparison, we applied Phenix on the same set of density maps. The dataset was aggregated by the EMDataResource and contained 62 high-resolution density maps, 52 of which have a deposited model PDB structure [39]. The dataset as well as the determined models will be actively updated at DeepTracer’s website as more and more data is deposited to EMDR. To our knowledge, this is the first CoV-related 3D cryo-EM modeling test dataset.

The scatter plots in Figure 11 show the evaluation results for the metrics calculated by Phenix’s `chain_comparison` tool, for the 52 coronavirus-related density maps that have a deposited model structure. The average percentage of matched model residues is 84% for DeepTracer and 49.8% for Phenix. This means that, on average, around 34% more residues were correctly placed by DeepTracer than by Phenix. The RMSD metric calculated an average value of 1.37Å for Phenix compared to 0.93Å with DeepTracer. Thus, DeepTracer not only determines more residues correctly than Phenix, but the correctly determined residues were also closer to the residues of the deposited model by around 0.4Å. For the sequence matching results, Phenix scored 24.95%, while Deep-Tracer achieved a sequence matching percentage of 63.08%. Finally, the mean length of consecutively matched residues in the modelled and deposited structure increased from 8.9 with Phenix to 20 with DeepTracer.

**Figure 11:**
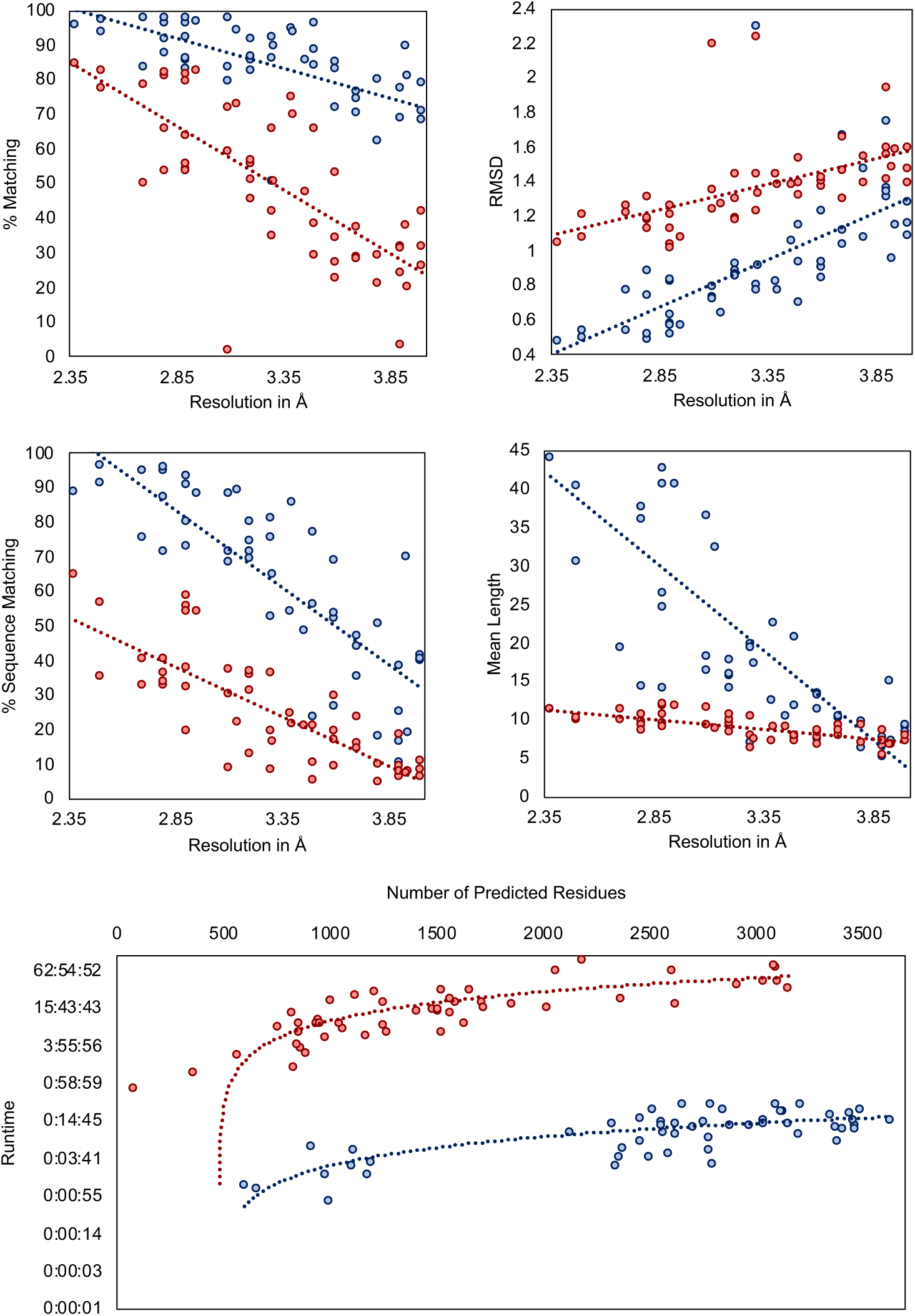
Results for coronavirus-related density maps. Evaluation of models built by DeepTracer (blue) and Phenix (red) for 52 coronavirus-related high-resolution density maps. The dotted lines represent the trend for each method. Computation times are shown on a logarithmic scale. Precise data can be found in Table S2.

The SARS-CoV-2 results from Table 1 show a similar pattern as the results of all coronavirus-related maps. DeepTracer outperformed Phenix in every metric with the most significant differences in the matching percentage and sequence matching. Additionally, the DeepTracer achieved a GDC score almost three times that of the Phenix method.

**Table 1:** Comparison of DeepTracer (DT) and Phenix (P) for SARS-CoV-2 dataset.

| EMDB | PDB | Residues | % Matching |  | RMSD |  | % Seq ID |  | GDC |  |
| --- | --- | --- | --- | --- | --- | --- | --- | --- | --- | --- |
|  |  |  | DT | P | DT | P | DT | P | DT | P |
| 21375 | 6vsb | 2905 | 84.90 | 48.60 | 1.14 | 1.40 | 45.90 | 20.90 | 17.88 | 5.39 |
| 21452 | 6vxx | 2916 | 91.40 | 53.80 | 0.96 | 1.18 | 61.30 | 40.00 | - | - |
| 30039 | 6m17 | 3072 | 80.30 | 53.10 | 1.72 | 1.72 | 69.80 | 54.60 | 11.84 | 8.31 |
| 30127 | 6m71 | 1077 | 91.70 | 54.20 | 1.02 | 1.20 | 58.60 | 16.60 | 20.74 | 8.89 |
| 30178 | 7btf | 1227 | 94.90 | 81.00 | 0.83 | 1.09 | 85.80 | 51.80 | 65.57 | 23.06 |
| 30209 | 7bv1 | 1102 | 87.60 | 67.00 | 0.84 | 1.29 | 87.50 | 30.50 | 55.62 | 18.15 |
| 30210 | 7bv2 | 1006 | 92.40 | 78.20 | 0.78 | 1.08 | 88.90 | 53.70 | 40.90 | 13.32 |
| Avg. |  |  | 89.03 | 62.27 | 1.04 | 1.28 | 71.11 | 38.30 | 35.42 | 12.85 |

GDC score could not be calculated for the EMD-21452 density map as the LGA web service could not process the modelled structures due to their size.

In Figure 12, we can see the structures modelled by DeepTracer for the EMD-30044 density map, which captures the human receptor angiotensin-converting enzyme 2 (ACE2) to which the spike protein of the SARS-CoV-2 virus binds to [11] and the EMD-21374 density map of a SARS-CoV-2 spike glycoprotein. No model structure has been deposited to the EMDR for either density map as of the date this paper is announced. This represents an ideal opportunity to showcase the potential of Deep-Tracer. Without any other parameters or manual processing steps, DeepTracer can determine detailed models based on the density maps. Researchers can use these models to develop therapeutics targeting the binding process between the spike protein and the human enzyme.

**Figure 12:**
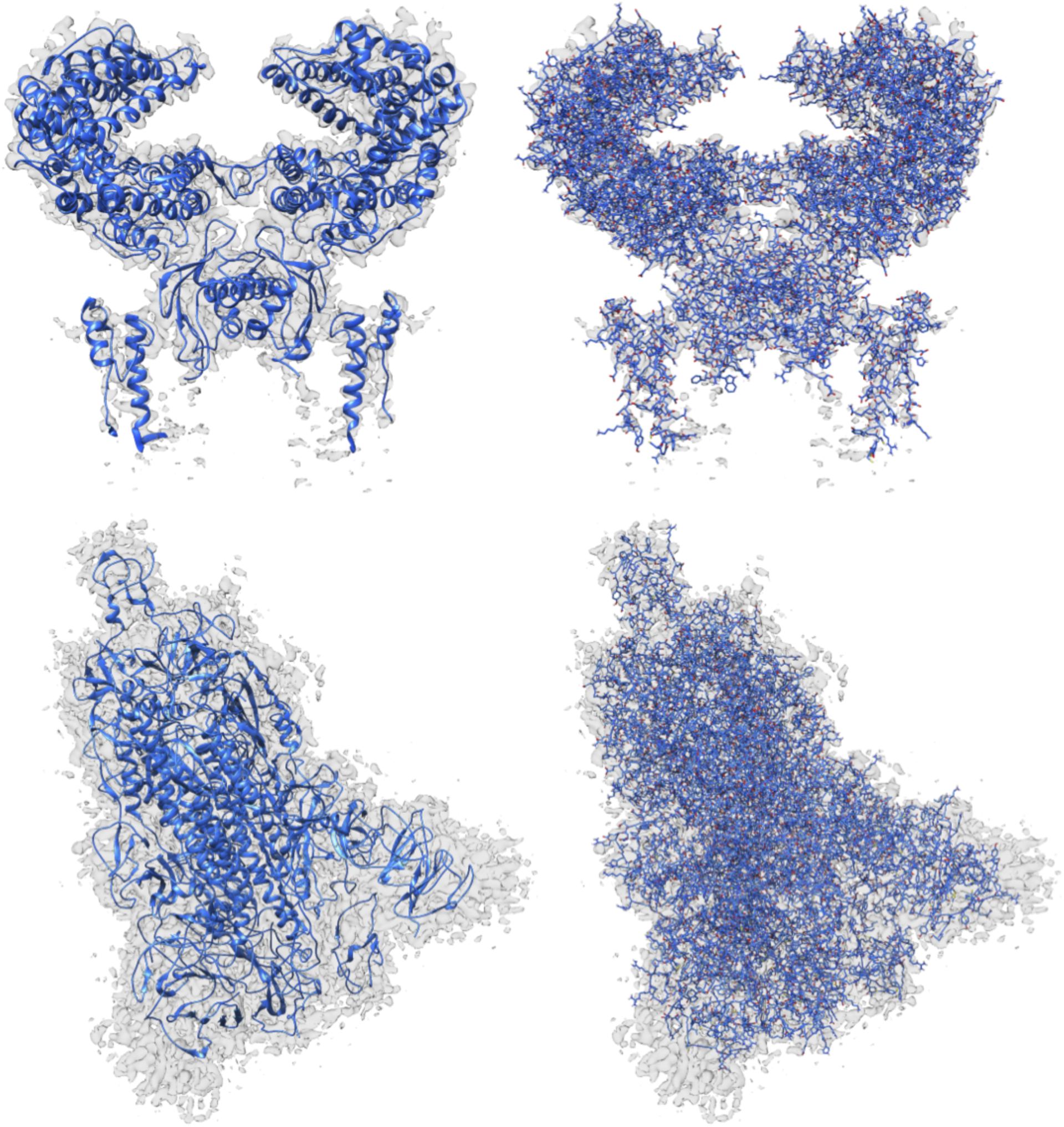
Models built from SARS-CoV-2 density maps, which do not have deposited model structures in the EMDR. DeepTracer model for the EMD-30044 density map (top) showing a human receptor angiotensin-converting enzyme 2 (ACE2) to which spike proteins of the SARS-CoV-2 virus bind to and the EMD-21374 depicting a SARS-CoV-2 spike glycoprotein. No model structure has been deposited to the EMDataResource for the density maps as of the date this paper is announced.

### 3.4 Computation Time

A major bottleneck of existing methods is their computational complexity, which renders them unable to model larger protein complexes. Thus, we conducted an analysis of DeepTracer’s computational time versus Phenix’s. The result is shown in Figure 11. The tests were executed on a machine with an Nvidia GeForce GTX 1080 Ti GPU, 8 processors, and 62 GB of memory. Although a comparison with the Phenix method is not entirely fair as Phenix does not take advantage of the machine’s GPU, this comparison provides a glimpse of the possibility that DeepTracer can achieve. We observed that Phenix took about 45 minutes to process a map containing 79 residues, while DeepTracer processed a map containing 2798 residues in only 26 minutes. Furthermore, the largest cryo-EM map (EMD-9891) that DeepTracer was tested on required around 14 minutes to complete, whereas Phenix’s processing time for this map was over 60 hours. DeepTracer is able to exploit the processing power of the GPU, which is becoming a staple on modern computing systems, to increase the throughput of scientific discovery. DeepTracer can model even very large protein complexes in a matter of hours. As an example, it traced around 60,000 residues for the EMD-9829 density map within only two hours.

## 4 Discussion

In this paper, we present DeepTracer, a fully-automatic tool that determines the allatom structures of protein complexes based on their cryo-EM density maps, using a tailored deep convolutional neural network and a set of computational methods. We applied this novel software on a set of coronavirus-related density maps and compared the results to Phenix, the state of the art cryo-EM model determination method [14]. We found that DeepTracer correctly placed, on average, around 30% more residues than Phenix with an average RMSD improvement of 0.11Å, from 1.29Å to 1.18Å. We also applied DeepTracer on a dataset of coronavirus-related density maps and calculated a coverage of 84% compared to 49.8% with Phenix and an average RMSD value of 0.93Å for DeepTracer and 1.37Å for Phenix. Detailed description and discussion can be found in the supplementary material. Furthermore, we compared DeepTracer with Rosetta and MAINMAST on a previously published set of nine density maps and observed significant RMSD improvements in comparison with Rosetta from 1.37Å to 0.85Å and much more complete model compared to MAINMAST with a coverage increase of 57%, from 36.4% to 93.4%. Detailed description and discussion can be found in the supplementary material. These results represent a significant accuracy boost, resulting in more complete protein structures. Particularly, for large protein complexes, DeepTracer built models much faster than other methods, tracing tens of thousands of residues with million of atoms within only a few hours. We achieved the results without any manual pre-processing steps, such as zoning or cutting of the density map using a deposited model structure. This means we can determine models without any prior knowledge about the cryo-EM map, and the users do not need to tune any parameters in order to obtain an accurate structure.

As the cryo-EM technology becomes more readily available, the number of captured density maps, especially larger protein complexes, is rising rapidly. DeepTracer allows for a greater throughput of cryo-EM as it can automatically and accurately infer structural information from density maps of macromolecule. This outcome ultimately accelerates the scientific discovery process, which is particularly urgent today, given the ongoing coronavirus pandemic. Coronavirus-related density maps are deposited to the EMDR on a daily basis. Our efficient and automated method to model these maps is an important tool for researchers to resolve the structural information of the virus-related macromolecules.

## Acknowledgements

This research was funded by the National Science Foundation (award #2030381) and the graduate research award of Computing and Software Systems division at University of Washington Bothell. We thank Yinrui (Bobby) Deng and other DAIS group members for the frontend development and testing of DeepTracer.

