## Supplementary Material for "DeepTracer: Fast Cryo-EM Protein Structure Modeling and Special Studies on CoV-related Complexes"

### Supplementary Materials of DeepTracer: Fasts Cryo-EM Protein Structure Modeling and Special Studies on CoV-related Complexes

Jonas Pfab[1], Nhut Minh Phan[1], and Dong Si[1, *]

[1]Division of Computing and Software Systems, University of Washington, Bothell, WA, 98011, USA

July 28, 2020

### S1 Pre-Processing

The goal of the pre-processing steps is to prepare the cryo-EM density maps for the neural network. These steps are crucial as maps can differ significantly in terms of shape, quality, resolution, and more. Therefore, we need to process the maps and convert them into a consistent format such that the neural network can understand connections across density maps. The pre-processing steps that achieve this are examined in Sections S1.1, S1.2, and S1.3.

#### S1.1 Data Grid Resampling

The first pre-processing step is to standardize the voxel size of all density maps. The grid storing the volume data of the density map has an associated voxel size, which determines the size of a single grid element or voxel in Angstrom. Without standardizing this voxel size to a fixed value the neural network could not draw conclusions about how far two voxels are from each other, making it difficult to predict the location of any amino acids. Therefore, this step ensures that each density maps has a voxel size of exactly 0.5Å. The value 0.5 was chosen based on several rounds of testing as a trade-off between prediction precision and memory usage of the resulting grids.

To set the voxel size of a density map to 0.5Å, we cannot simply change the meta data of the map. We had to resample the volume data onto a new grid in which each voxel represents 0.5Å. An example of a resampling process from an origin grid to a grid with half the voxel size is shown in Figure S1. Here, the shape of the volume remains the same, however, we require eight times the number of voxels to represent it. To realize the resampling step, DeepTracer utilizes UCSF Chimera (Eric F Pettersen et al. Ucsf chimera — a visualization system for exploratory research and analysis. Journal of computational chemistry, 25(13):1605–1612, 2004). First, it creates a new grid of the same size in Angstrom as the original density map, but with a voxel size of 0.5Å. Then, it uses Chimera's resampling command (vop. `https://www.cgl.ucsf.edu/chimera/docs/UsersGuide/midas/vop.html#resample`. (Accessed on 03/01/2020)) to resample the original density map onto the newly created grid.

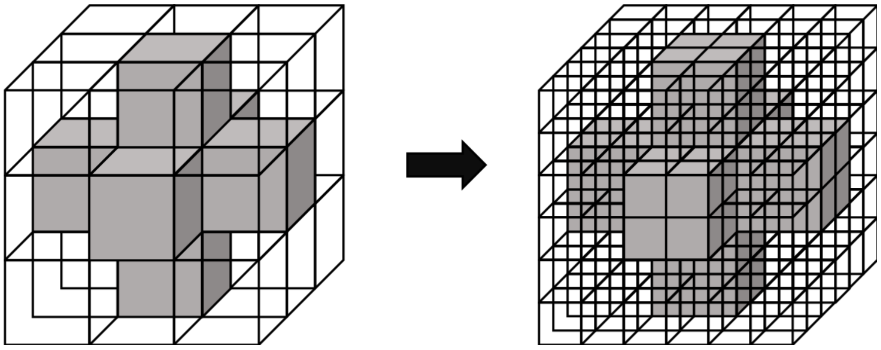

Figure S1: Data grid resampling. Visualization of the resampling process from an origin grid onto a grid with half the voxel size.

#### S1.2 Density Value Normalization

The absolute value of a voxel in itself contains little information. We have to rely on the density values of other voxels to make conclusions about the protein structure. Consequently, we can normalize the density values without risking information loss. The normalization process makes sure that the range of density values is identical for all density maps. In the case of experimental maps, this range can initially differ substantially with some maps contain values from -0.1 to 0.1 and other maps ranging from -10 to 20.

To normalize values, we can usually divide each value by the overall highest value. However, this process is problematic for some density maps as there are outlier density values, which have values that are much higher than all other values. If we were to divide all other values using these outlier, all other density values would end up being close to zero. Therefore, we used the 95th percentile of the density values to divide all other values with. Afterwards, we simply set the few values that are greater than 1 to 1. Additionally, we set all values below 0 to 0 as they contain no valuable information for our use case. This leaves us with a range from 0 to 1, which contains all density values. An example of the density value histograms before and after the normalization step can be seen in Figure S2.

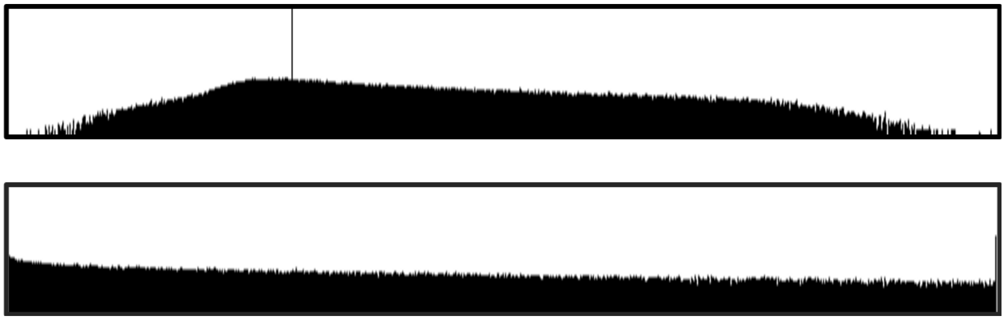

Figure S2: Density value normalization. Histograms of the 6272 density map depicting the relative frequency of density values before (top) and after (bottom) normalization. The vertical black line in the upper histogram indicates the value 0 and is not visible in the lower one as there are no density values below 0 anymore.

#### S1.3 Grid Division

The input layer of the model takes a density map to make a prediction. We have to make sure that the dimensions of the volume data grid of the density map are identical to those of the input layer of the model to avoid mismatching errors. However, the dimensions of the grid vary from map to map, demanding modification of the grid to match its dimensions of the input layer. Unfortunately, we cannot simply scale the density map for it to fit the input layer as this would change the size of each voxel in Angstrom, which has to remain 0.5 as mentioned in Section S1.1. Therefore, we divided the grid into multiple sub grids each the size of the input layer of the deep learning model.

We divided the volume data grid into sub grids of size $64^3$. The number 64 was chosen as it creates a relatively small input layer that is still broad enough for the deep learning model to detect larger patterns, such as secondary structure elements. Dividing the grid, however, can aggravate predictions in areas close to the border of sub grids as relevant information from neighboring voxels might be cut off. Therefore, we introduce a core grid of size $50^3$ in the center of each sub grid. Although each sub grid has a size of $64^3$, we only used the predictions from the inner core grid. Consequently, when dividing the grid we overlapped the $64^3$ sub grids such that core grids of all sub grids cover the entire original grid without overlap. An example of such a division for a two-dimensional grid can be seen in Figure S3.

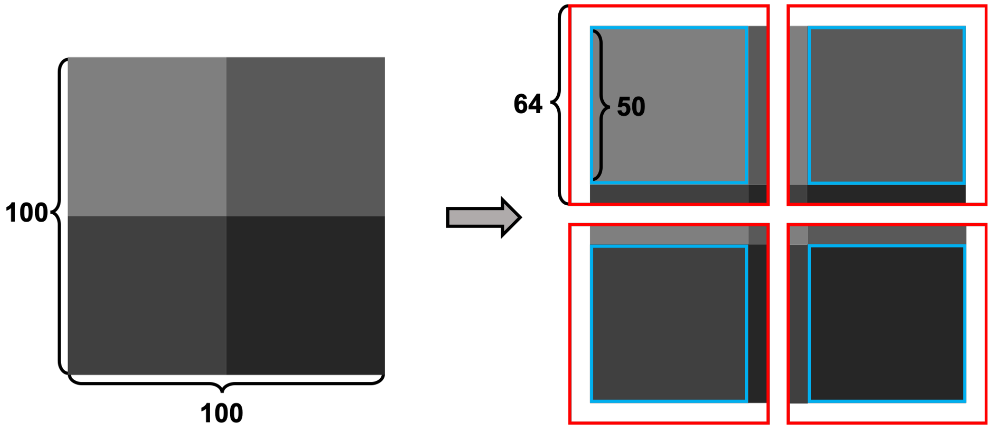


Figure S3: Grid division into sub-grids. Visualization of the division of a $100^2$ grid into four grids each with a dimension of $64^2$ and a core size of $50^2$. The red lines indicate the complete sub grids while the blue lines show their cores.

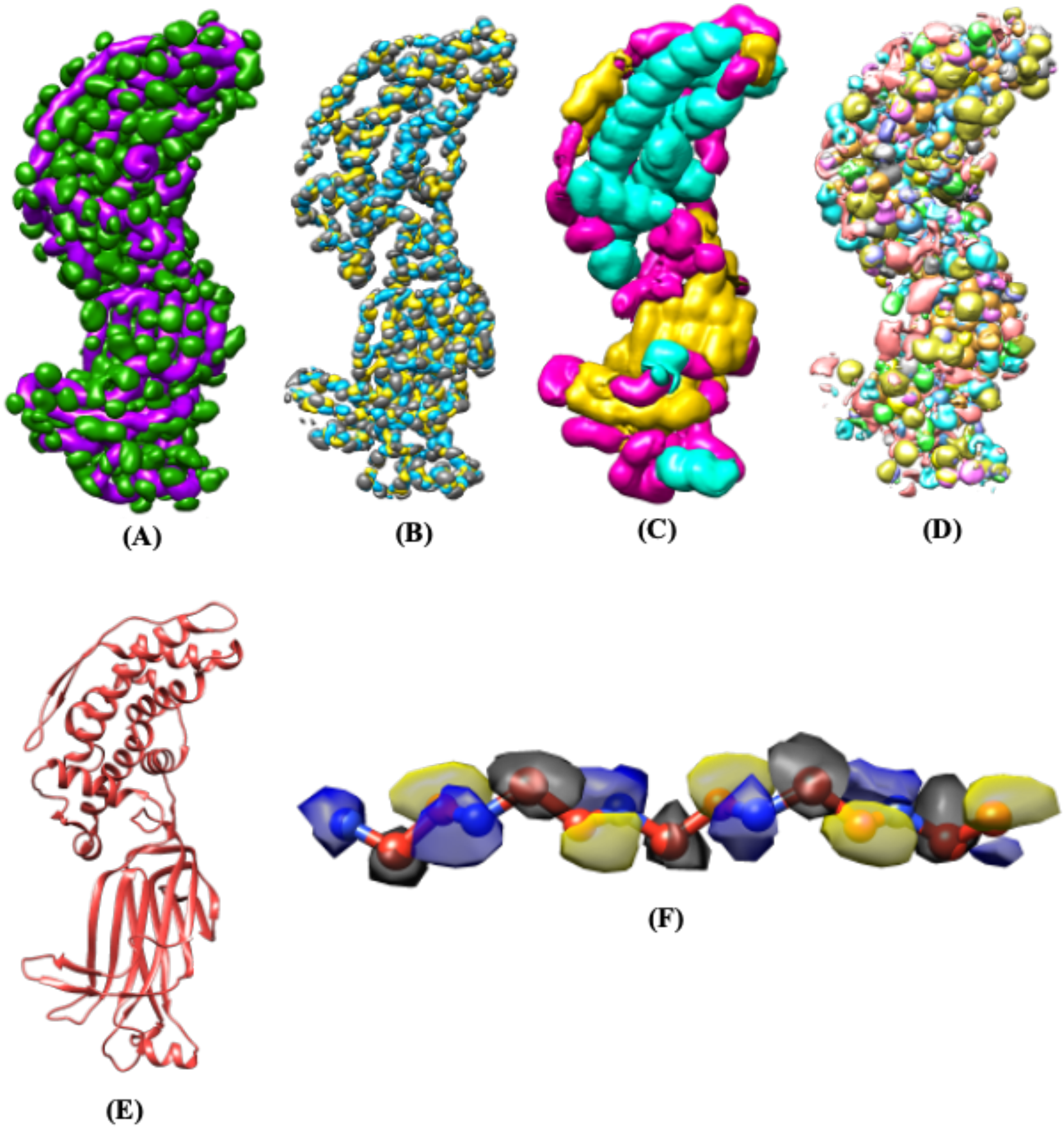


Figure S4: Neural network predictions. Raw prediction of the U-Net deep learning model for the 6272 density map. (A) Backbone prediction with backbone in purple and side-chains in green. (B) Atoms prediction with Cα atoms in gray, C atoms in yellow, and N atoms in blue. (C) Secondary structure prediction with α-helices in turquoise, β-sheets in yellow and loops in pink. (D) Amino acid type prediction with a different color for each type. (E) Solved structure (PDB-3J9S) of density map. (F) Atoms prediction segment next to solved structure.

### S2 Comparison with MAINMAST and Rosetta

In addition to Phenix, MAINMAST and Rosetta are two further established cryo-EM prediction methods. We conducted a brief analysis of their performances compared to DeepTracer based on a test set of nine density maps taken from the previous papers (Frenz B et al., Nature Methods, 14:797-800, 2017), (Terashi, G., Kihara, D. De novo main-chain modeling for EM maps using MAINMAST. Nat Commun 9, 1618 (2018)). Note that the density maps were cropped such that they captured only a single protein chain. This cropping was necessary as both methods can only perform single-chain predictions. To evaluate predictions we utilize Phenix's `chain_comparison` tool. The results of this analysis can be seen in Table S1. We can note that DeepTracer outperforms Rosetta in all four metrics with particularly significant improvements in the percentage of matched residues as well as false-positive predictions. Compared to the MAINMAST method DeepTracer performed worse in three of the four metrics. However, predictions of DeepTracer were much more complete with an average matching percentage of 93.4% compared to only 36.4% with MAINMAST. That means that MAINMAST correctly predicted only around 1/3 of the protein structure.

Table S1: Comparison of DeepTracer with MAINMAST and Rosetta on a dataset of 9 density maps.

| Method | Protein | % Matching | RMSD | % Seq Matching | % FP |
|---|---|---|---|---|---|
| DeepTracer | BPP1 | 93.30 | 0.82 | 66.20 | 2.56 |
| | FrhB | 98.20 | 0.67 | 68.40 | 3.17 |
| | T20S | 86.40 | 1.19 | 33.50 | 4.55 |
| | VP6 | 93.20 | 0.98 | 48.10 | 2.38 |
| | TRPV1 | 87.70 | 0.85 | 71.30 | 3.20 |
| | FrhA | 97.90 | 0.60 | 98.10 | 1.31 |
| | FrhG | 96.10 | 0.86 | 90.40 | 5.63 |
| | STIV | 90.40 | 0.88 | 70.10 | 0.32 |
| | TMV | 97.40 | 0.84 | 58.30 | 3.21 |
| | **Avg.** | **93.40** | **0.85** | **67.16** | **2.93** |
| Rosetta | BPP1 | 73.10 | 1.51 | 58.20 | 28.44 |
| | FrhB | 90.00 | 1.07 | 93.30 | 10.68 |
| | T20S | 77.40 | 1.61 | 62.60 | 24.89 |
| | VP6 | 72.00 | 1.37 | 48.60 | 28.72 |
| | TRPV1 | 84.20 | 1.22 | 69.70 | 18.10 |
| | FrhA | 92.50 | 0.90 | 93.30 | 8.29 |
| | FrhG | 63.20 | 1.79 | 37.50 | 41.23 |
| | STIV | 47.70 | 1.65 | 46.30 | 52.75 |
| | TMV | 91.00 | 1.20 | 88.70 | 10.97 |
| | **Avg.** | **76.79** | **1.37** | **66.47** | **24.89** |
| MAINMAST | BPP1 | 17.40 | 0.68 | 100.00 | 0.00 |
| | FrhB | 59.60 | 0.77 | 98.80 | 2.34 |
| | T20S | 23.10 | 1.01 | 100.00 | 1.96 |
| | VP6 | 26.20 | 0.73 | 99.00 | 0.00 |
| | TRPV1 | 33.20 | 0.72 | 99.00 | 0.00 |
| | FrhA | 57.70 | 0.62 | 97.70 | 0.89 |
| | FrhG | 31.10 | 0.72 | 100.00 | 1.39 |
| | STIV | 14.80 | 0.68 | 100.00 | 0.00 |
| | TMV | 64.50 | 0.78 | 98.00 | 0.99 |
| | **Avg.** | **36.40** | **0.75** | **99.17** | **0.84** |

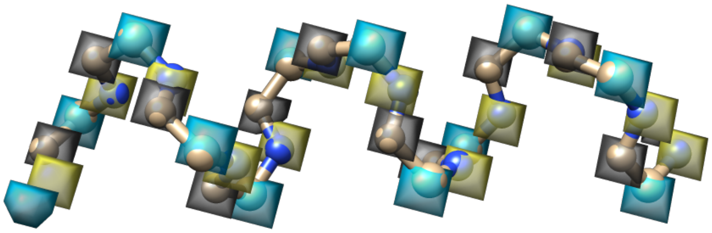

Figure S5: Atom mask. Portion of the atom mask containing backbone atoms for part of a helix from the PDB-6NQ1 structure. The gray labels indicate carbon alpha atoms, the blue labels carbon atoms, and the yellow labels nitrogen atoms.

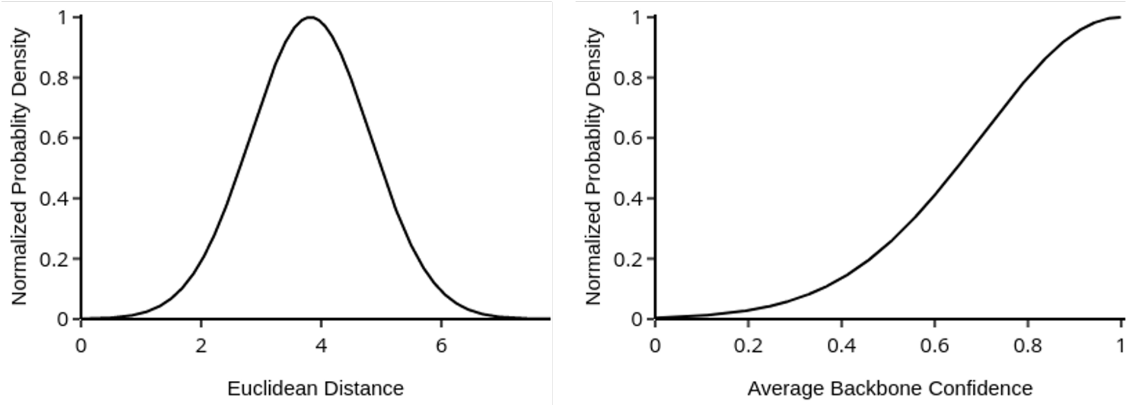


Figure S6: Probability density function for connection confidence. Normalized probability density function used to calculate confidence score for the euclidean distance and average backbone confidence between two Cα atoms. This is used to express a distance between two atoms for the traveling salesman algorithm tracing the backbone.

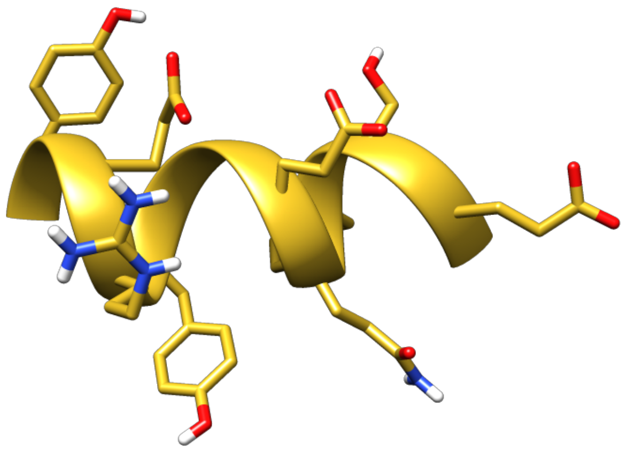

Figure S7: Predicted side chains for an extract of an α-helix. Backbone atoms are displayed in the ribbon view and side chain atoms in atom view.

Table S2: Precise prediction results for coronavirus-related maps. Evaluation of prediction results from DeepTracer and Phenix for 52 coronavirus-related high-resolution density maps.

| | **DeepTracer** | | | | **Phenix** | | | |
|---|---|---|---|---|---|---|---|---|
| EMDB ID | % Matching | RMSD | % Seq Matching | Mean Length | % Matching | RMSD | % Seq Matching | Mean Length |
| 0401 | 84.30 | 0.93 | 55.90 | 11.70 | 29.00 | 1.39 | 9.90 | 7.70 |
| 0402 | 83.10 | 0.90 | 51.80 | 13.40 | 34.40 | 1.40 | 19.30 | 7.60 |
| 0520 | 97.70 | 0.73 | 87.70 | 36.60 | 71.80 | 1.24 | 37.10 | 9.20 |
| 0521 | 88.80 | 1.15 | 23.30 | 7.70 | 38.20 | 1.53 | 5.10 | 7.20 |
| 0557 | 96.10 | 0.48 | 94.60 | 37.60 | 80.90 | 1.13 | 39.90 | 9.70 |
| 10676 | 94.60 | 0.82 | 53.80 | 12.60 | 75.00 | 1.44 | 24.50 | 7.30 |
| 10863 | 31.40 | 1.74 | 10.10 | 5.10 | 3.50 | 1.94 | 18.20 | 5.50 |
| 11007 | 85.60 | 0.58 | 93.10 | 42.70 | 63.90 | 1.03 | 58.40 | 11.90 |
| 20070 | 96.10 | 0.52 | 92.80 | 49.10 | 81.50 | 1.21 | 37.60 | 9.60 |
| 20542 | 97.40 | 0.54 | 95.80 | 40.30 | 82.60 | 1.21 | 34.80 | 9.90 |
| 20543 | 97.80 | 0.52 | 95.60 | 47.20 | 82.20 | 1.18 | 35.90 | 10.60 |
| 20544 | 97.90 | 0.54 | 94.50 | 49.30 | 78.30 | 1.22 | 40.10 | 10.00 |
| 20545 | 97.80 | 0.57 | 90.30 | 40.50 | 79.60 | 1.26 | 31.90 | 9.60 |
| 20668 | 79.50 | 0.72 | 67.90 | 18.30 | 59.10 | 1.35 | 29.80 | 9.20 |
| 20672 | 94.00 | 0.64 | 88.90 | 32.30 | 72.70 | 1.27 | 21.60 | 8.80 |
| 21375 | 85.60 | 1.05 | 48.30 | 10.50 | 47.60 | 1.38 | 20.80 | 9.10 |
| 21377 | 80.80 | 1.15 | 18.90 | 7.20 | 20.30 | 1.58 | 7.60 | 6.70 |
| 21391 | 96.20 | 0.70 | 76.70 | 20.70 | 65.70 | 1.32 | 20.90 | 7.90 |
| 21452 | 91.60 | 0.88 | 70.90 | 14.40 | 53.70 | 1.19 | 32.60 | 9.30 |
| 21457 | 91.80 | 0.88 | 71.20 | 15.70 | 55.60 | 1.19 | 31.20 | 9.50 |
| 21864 | 83.40 | 0.79 | 70.90 | 16.40 | 1.90 | 2.19 | 8.70 | 11.50 |
| 21865 | 76.60 | 1.03 | 43.70 | 10.20 | 28.60 | 1.30 | 15.60 | 9.30 |
| 21997 | 83.50 | 0.77 | 75.10 | 19.30 | 50.00 | 1.26 | 32.30 | 11.30 |
| 21999 | 85.90 | 0.80 | 75.30 | 19.80 | 41.90 | 1.23 | 19.30 | 10.40 |
| 22000 | 72.00 | 0.84 | 53.10 | 11.30 | 27.20 | 1.37 | 16.50 | 8.40 |
| 22001 | 85.50 | 0.92 | 74.10 | 16.00 | 51.30 | 1.30 | 35.50 | 10.00 |
| 30037 | 96.00 | 0.47 | 88.60 | 44.10 | 84.80 | 1.04 | 64.50 | 11.40 |
| 30039 | 83.20 | 0.82 | 79.60 | 24.60 | 54.20 | 1.12 | 55.50 | 10.70 |
| 30040 | 86.00 | 0.63 | 72.40 | 26.50 | 55.70 | 1.01 | 53.80 | 12.00 |
| 30127 | 92.10 | 0.83 | 53.60 | 14.10 | 53.60 | 1.21 | 19.40 | 9.10 |
| 30178 | 97.00 | 0.57 | 87.90 | 40.60 | 82.50 | 1.07 | 53.90 | 11.80 |
| 30209 | 87.60 | 0.74 | 86.60 | 36.10 | 66.00 | 1.31 | 33.40 | 8.70 |
| 30210 | 93.60 | 0.49 | 90.70 | 30.50 | 77.70 | 1.07 | 56.50 | 10.10 |
| 6526 | 70.90 | 1.16 | 39.40 | 9.00 | 31.50 | 1.39 | 8.00 | 7.30 |
| 6703 | 82.30 | 0.86 | 69.10 | 14.20 | 45.40 | 1.44 | 12.80 | 8.40 |
| 6704 | 70.60 | 1.66 | 35.20 | 7.80 | 28.00 | 1.65 | 23.40 | 8.00 |
| 6705 | 74.50 | 1.11 | 46.60 | 10.40 | 37.10 | 1.46 | 14.20 | 8.60 |
| 6732 | 79.90 | 1.07 | 50.10 | 9.80 | 29.30 | 1.39 | 9.40 | 9.30 |
| 7063 | 92.20 | 0.77 | 81.00 | 19.40 | 66.00 | 1.44 | 8.00 | 7.80 |
| 7573 | 86.60 | 0.85 | 79.90 | 17.70 | 56.50 | 1.18 | 36.30 | 10.60 |
| 7574 | 69.00 | 1.36 | 24.70 | 7.20 | 31.60 | 1.55 | 6.20 | 6.50 |
| 7575 | 68.80 | 1.31 | 38.10 | 7.70 | 24.20 | 1.41 | 9.30 | 8.60 |
| 7577 | 50.60 | 2.29 | 52.40 | 6.90 | 34.70 | 2.23 | 35.80 | 6.20 |
| 7578 | 62.10 | 1.47 | 17.60 | 6.20 | 20.90 | 1.54 | 4.50 | 7.00 |
| 7631 | 89.90 | 0.95 | 69.80 | 14.90 | 37.60 | 1.48 | 7.00 | 6.70 |
| 8331 | 93.80 | 0.77 | 85.50 | 22.50 | 69.60 | 1.38 | 21.50 | 9.10 |
| 8783 | 68.40 | 1.28 | 41.30 | 8.30 | 26.30 | 1.59 | 10.80 | 8.00 |
| 8784 | 85.10 | 0.93 | 68.70 | 13.20 | 53.10 | 1.40 | 29.30 | 8.60 |
| 8791 | 79.20 | 1.08 | 39.90 | 9.30 | 41.90 | 1.47 | 6.20 | 7.80 |
| 9588 | 85.10 | 1.23 | 26.60 | 7.50 | 22.80 | 1.42 | 9.10 | 6.80 |
| 9589 | 77.30 | 1.34 | 16.40 | 6.60 | 31.30 | 1.59 | 7.50 | 7.20 |
| 9891 | 89.80 | 0.91 | 64.40 | 17.20 | 50.30 | 1.33 | 16.10 | 7.50 |

Table S3: Results of DeepTracer and Phenix for dataset of 476 density maps. Evaluation was accomplished using Phenix's `chain_comparison` tool. For three rows the tool did not return any values for Phenix's prediction and the corresponding cells are filled with a '-'.

| | **DeepTracer** | | | | **Phenix** | | | |
|---|---|---|---|---|---|---|---|---|
| EMDB ID | % Matching | RMSD | % Seq Matching | Mean Length | % Matching | RMSD | % Seq Matching | Mean Length |
| 2278 | 87.60 | 1.31 | 64.20 | 12.70 | 64.50 | 1.24 | 17.10 | 8.10 |
| 2364 | 59.90 | 1.80 | 5.70 | 4.90 | 49.40 | 1.91 | 9.00 | 5.40 |
| 2513 | 98.10 | 0.81 | 90.30 | 26.50 | 62.60 | 1.17 | 11.80 | 6.60 |
| 2566 | 95.20 | 0.79 | 84.90 | 23.00 | 51.60 | 0.99 | 21.00 | 9.10 |
| 2599 | 77.10 | 2.05 | 44.20 | 7.90 | 22.80 | 1.96 | 9.50 | 6.30 |
| 2650 | 92.30 | 0.82 | 90.40 | 28.30 | 59.80 | 0.98 | 19.40 | 9.30 |
| 2699 | 63.30 | 1.22 | 60.30 | 15.80 | 3.50 | 1.20 | 30.30 | 10.50 |
| 2762 | 92.10 | 0.88 | 73.00 | 17.80 | 45.80 | 1.09 | 15.00 | 8.10 |
| 2763 | 65.50 | 1.23 | 19.30 | 7.10 | 26.30 | 1.33 | 10.60 | 7.60 |
| 2764 | 76.00 | 1.19 | 28.40 | 8.30 | 29.50 | 1.22 | 9.00 | 7.30 |
| 2773 | 74.10 | 1.43 | 19.80 | 7.20 | 12.60 | 1.40 | 8.50 | 6.60 |
| 2807 | 73.00 | 1.38 | 48.90 | 9.80 | 44.20 | 1.34 | 7.60 | 7.20 |
| 2832 | 88.70 | 1.19 | 78.10 | 20.10 | 32.20 | 1.33 | 7.80 | 8.00 |
| 2842 | 93.50 | 0.89 | 96.50 | 20.40 | 60.80 | 1.08 | 23.70 | 8.50 |
| 2847 | 85.20 | 0.92 | 65.70 | 12.90 | 20.30 | 0.84 | 17.60 | 9.20 |
| 2850 | 86.80 | 1.13 | 36.20 | 8.40 | 63.50 | 1.39 | 5.90 | 6.10 |
| 2856 | 77.70 | 1.10 | 55.70 | 10.70 | 49.70 | 1.38 | 7.60 | 6.80 |
| 2857 | 72.60 | 1.09 | 65.40 | 10.90 | 45.30 | 1.22 | 8.20 | 6.90 |
| 2867 | 88.30 | 1.36 | 67.90 | 9.80 | 68.70 | 1.50 | 8.50 | 7.40 |
| 2870 | 52.30 | 1.58 | 34.80 | 7.90 | 30.60 | 1.36 | 4.50 | 7.60 |
| 2871 | 50.80 | 1.59 | 46.70 | 9.60 | 38.40 | 1.70 | 2.70 | 7.70 |
| 2913 | 80.90 | 0.95 | 46.60 | 12.10 | 33.50 | 1.10 | 9.10 | 7.70 |
| 2924 | 78.90 | 1.10 | 66.30 | 14.50 | 56.70 | 0.97 | 10.80 | 10.30 |
| 2974 | 69.40 | 1.37 | 11.00 | 8.20 | 52.60 | 1.48 | 6.90 | 7.20 |
| 2981 | 84.30 | 1.23 | 13.30 | 7.00 | 62.90 | 1.28 | 9.80 | 7.40 |
| 2984 | 96.30 | 0.63 | 91.40 | 36.40 | 87.90 | 0.59 | 56.30 | 18.00 |
| 3013 | 97.80 | 0.56 | 98.20 | 68.10 | 68.90 | 1.02 | 7.00 | 7.10 |
| 3014 | 98.20 | 0.54 | 97.10 | 55.30 | 75.70 | 0.74 | 39.20 | 11.50 |
| 3019 | 63.80 | 0.87 | 63.80 | 14.00 | 26.10 | 1.11 | 8.80 | 7.60 |
| 3047 | 84.50 | 0.98 | 63.20 | 13.20 | 38.00 | 1.10 | 15.40 | 8.00 |
| 3061 | 94.90 | 0.89 | 92.30 | 25.20 | 73.90 | 1.15 | 12.10 | 9.50 |
| 3062 | 33.90 | 1.75 | 14.00 | 6.50 | 61.20 | 1.50 | 6.20 | 6.80 |
| 3063 | 47.40 | 1.53 | 10.90 | 7.50 | 66.50 | 1.52 | 5.80 | 6.20 |
| 3121 | 73.90 | 1.28 | 37.10 | 7.30 | 56.30 | 1.63 | 5.20 | 5.60 |
| 3129 | 94.40 | 0.61 | 95.00 | 56.50 | 78.90 | 0.94 | 25.80 | 10.80 |
| 3130 | 88.60 | 0.66 | 91.00 | 39.30 | 71.30 | 1.20 | 16.70 | 8.00 |
| 3133 | 47.50 | 1.64 | 11.20 | 5.90 | 9.70 | 1.64 | 10.30 | 6.80 |
| 3137 | 82.20 | 1.62 | 30.10 | 6.50 | 56.70 | 1.74 | 7.60 | 5.40 |
| 3151 | 87.80 | 0.83 | 71.00 | 17.60 | 34.20 | 0.98 | 11.20 | 8.00 |
| 3152 | 77.30 | 1.05 | 24.90 | 8.20 | 24.30 | 1.16 | 9.60 | 7.60 |
| 3178 | 83.20 | 1.30 | 37.10 | 9.70 | 44.90 | 1.37 | 9.30 | 7.10 |
| 3218 | 88.40 | 0.80 | 87.50 | 23.90 | 49.90 | 1.10 | 13.20 | 7.80 |
| 3222 | 85.30 | 1.49 | 42.40 | 7.30 | 29.40 | 1.74 | 4.20 | 4.80 |
| 3231 | 91.30 | 1.05 | 19.20 | 10.60 | 69.10 | 1.18 | 6.90 | 8.00 |
| 3236 | 87.60 | 1.54 | 59.50 | 9.50 | 11.10 | 1.67 | 29.20 | 12.00 |
| 3237 | 85.00 | 1.38 | 12.90 | 8.60 | 64.20 | 1.47 | 9.90 | 7.90 |
| 3238 | 70.60 | 1.40 | 15.60 | 7.90 | 56.10 | 1.50 | 7.30 | 8.00 |
| 3239 | 82.00 | 1.24 | 18.90 | 9.50 | 63.00 | 1.34 | 7.40 | 8.50 |
| 3240 | 69.80 | 1.49 | 13.80 | 8.10 | 68.30 | 1.54 | 8.50 | 8.00 |
| 3246 | 90.90 | 1.09 | 36.60 | 12.30 | 62.60 | 1.47 | 9.10 | 6.70 |
| 3295 | 77.20 | 0.96 | 77.10 | 13.60 | 64.20 | 0.97 | 32.10 | 9.90 |
| 3296 | 79.50 | 0.96 | 78.40 | 12.00 | 67.40 | 0.93 | 44.40 | 10.80 |
| 3297 | 88.90 | 1.11 | 71.40 | 12.10 | 57.80 | 1.01 | 16.00 | 8.40 |
| 3298 | 87.50 | 1.10 | 73.50 | 12.00 | 60.40 | 0.94 | 20.70 | 8.30 |
| 3299 | 77.20 | 1.09 | 75.70 | 14.20 | 61.50 | 1.02 | 19.00 | 8.50 |

| | | | | | | | | |
|---|---|---|---|---|---|---|---|---|
| 3308 | 85.20 | 1.12 | 37.50 | 8.20 | 52.90 | 1.38 | 5.20 | 6.60 |
| 3337 | 77.50 | 1.32 | 20.20 | 8.50 | 49.90 | 1.39 | 5.40 | 6.40 |
| 3366 | 53.20 | 1.60 | 10.80 | 6.40 | 41.80 | 1.73 | 5.50 | 5.80 |
| 3378 | 50.10 | 1.37 | 14.80 | 6.70 | 37.00 | 1.47 | 7.50 | 6.50 |
| 3385 | 76.90 | 1.31 | 34.90 | 10.50 | 51.00 | 1.26 | 10.80 | 9.60 |
| 3388 | 80.90 | 1.12 | 63.80 | 14.30 | 54.90 | 1.00 | 12.50 | 9.80 |
| 3446 | 82.10 | 1.16 | 44.80 | 11.20 | 50.20 | 1.43 | 6.60 | 6.80 |
| 3447 | 70.70 | 1.50 | 10.20 | 6.10 | 31.40 | 1.64 | 7.60 | 5.80 |
| 3448 | 77.90 | 1.41 | 16.40 | 7.10 | 29.50 | 1.55 | 6.80 | 7.00 |
| 3460 | 71.20 | 1.23 | 45.30 | 10.30 | 46.80 | 1.28 | 12.60 | 7.60 |
| 3489 | 88.80 | 0.70 | 89.70 | 24.30 | 51.10 | 0.79 | 22.00 | 10.30 |
| 3490 | 86.20 | 0.69 | 87.20 | 22.10 | 38.90 | 0.83 | 14.20 | 9.70 |
| 3492 | 83.60 | 0.88 | 68.20 | 14.20 | 23.00 | 0.94 | 15.20 | 7.80 |
| 3493 | 85.90 | 0.69 | 83.20 | 20.30 | 36.00 | 0.85 | 26.80 | 9.90 |
| 3508 | 90.40 | 0.83 | 78.80 | 18.10 | 39.20 | 0.90 | 17.40 | 9.30 |
| 3523 | 62.30 | 1.70 | 8.80 | 6.40 | 54.50 | 1.79 | 7.70 | 6.20 |
| 3524 | 80.70 | 1.36 | 49.20 | 9.90 | 60.50 | 1.48 | 6.50 | 6.60 |
| 3525 | 81.70 | 1.04 | 35.80 | 9.80 | 26.50 | 1.14 | 13.70 | 7.30 |
| 3533 | 69.10 | 0.95 | 52.20 | 10.70 | 6.10 | 0.99 | 12.20 | 8.10 |
| 3538 | 96.10 | 1.34 | 98.60 | 37.00 | 85.10 | 1.56 | 10.70 | 7.70 |
| 3541 | 51.00 | 1.63 | 9.90 | 6.40 | 25.20 | 1.57 | 7.00 | 6.80 |
| 3570 | 87.90 | 0.91 | 32.10 | 8.70 | 59.80 | 1.38 | 6.00 | 6.40 |
| 3571 | 89.30 | 1.18 | 71.30 | 15.60 | 39.50 | 1.48 | 9.40 | 8.50 |
| 3574 | 89.60 | 0.71 | 88.30 | 24.00 | 56.20 | 0.94 | 29.40 | 9.90 |
| 3575 | 84.00 | 1.11 | 17.60 | 6.70 | 36.90 | 1.53 | 6.30 | 5.00 |
| 3583 | 91.20 | 1.13 | 68.40 | 13.60 | 67.20 | 1.18 | 9.30 | 10.80 |
| 3589 | 87.20 | 1.26 | 25.50 | 9.00 | 1.50 | 1.40 | 0.00 | 6.50 |
| 3593 | 70.30 | 1.01 | 75.30 | 16.80 | 46.40 | 1.27 | 14.10 | 7.50 |
| 3601 | 86.30 | 1.18 | 43.80 | 10.10 | 50.00 | 1.29 | 10.60 | 7.10 |
| 3618 | 81.80 | 1.00 | 55.70 | 11.30 | 2.90 | 1.03 | 8.50 | 12.60 |
| 3622 | 28.60 | 1.83 | 8.00 | 6.30 | 35.30 | 1.99 | 4.80 | 5.50 |
| 3624 | 68.10 | 1.27 | 31.20 | 8.10 | 5.80 | 1.24 | 14.20 | 7.10 |
| 3625 | 68.50 | 1.27 | 30.60 | 8.50 | 6.40 | 1.26 | 16.00 | 7.10 |
| 3630 | 96.60 | 0.71 | 95.70 | 35.20 | 76.50 | 0.87 | 32.50 | 8.40 |
| 3631 | 95.70 | 0.73 | 91.40 | 30.00 | 71.10 | 1.13 | 15.40 | 8.20 |
| 3652 | 57.90 | 1.11 | 67.50 | 27.90 | 48.20 | 1.07 | 24.90 | 10.00 |
| 3656 | 75.60 | 1.30 | 15.90 | 6.40 | 6.10 | 1.27 | 9.40 | 7.80 |
| 3695 | 66.40 | 1.38 | 19.90 | 7.20 | 31.80 | 1.47 | 9.40 | 6.60 |
| 3713 | 91.60 | 0.75 | 89.60 | 24.10 | 36.10 | 0.81 | 20.60 | 10.60 |
| 3727 | 30.60 | 1.63 | 10.30 | 6.10 | 33.00 | 1.78 | 8.30 | 6.10 |
| 3730 | 87.00 | 0.99 | 48.20 | 12.00 | 23.80 | 0.97 | 11.60 | 7.20 |
| 3741 | 91.80 | 0.80 | 62.70 | 11.20 | 63.00 | 0.92 | 10.90 | 11.50 |
| 3742 | 98.60 | 0.72 | 79.20 | 36.00 | 60.30 | 1.10 | 11.40 | 11.00 |
| 3743 | 91.80 | 0.96 | 59.70 | 11.20 | 4.10 | 1.53 | 0.00 | 3.00 |
| 3746 | 82.60 | 1.13 | 28.70 | 7.10 | 19.80 | 1.59 | 9.80 | 5.10 |
| 3748 | 85.70 | 0.86 | 68.90 | 13.40 | 25.10 | 1.06 | 13.70 | 8.00 |
| 3750 | 83.70 | 0.76 | 75.10 | 15.60 | 29.40 | 0.87 | 14.90 | 8.30 |
| 3766 | 18.80 | 1.70 | 9.00 | 6.80 | 17.40 | 1.78 | 6.40 | 6.90 |
| 3770 | 57.10 | 1.32 | 20.90 | 7.80 | 13.50 | 1.28 | 6.50 | 7.20 |
| 3785 | 90.30 | 1.29 | 79.70 | 20.80 | 42.50 | 1.74 | 12.50 | 6.30 |
| 3802 | 68.20 | 1.51 | 9.50 | 6.10 | 54.40 | 1.57 | 7.30 | 6.40 |
| 3817 | 71.40 | 0.88 | 77.30 | 16.40 | 43.50 | 1.18 | 11.00 | 8.40 |
| 3824 | 84.70 | 1.33 | 30.10 | 10.10 | 63.90 | 1.23 | 6.40 | 10.40 |
| 3842 | 64.80 | 1.31 | 19.50 | 6.60 | 28.30 | 1.63 | 6.00 | 6.20 |
| 3843 | 61.90 | 1.28 | 19.20 | 6.60 | 30.80 | 1.68 | 9.90 | 5.90 |
| 3847 | 70.60 | 1.05 | 41.40 | 10.50 | 34.50 | 1.18 | 8.80 | 7.40 |
| 3851 | 88.10 | 1.11 | 37.80 | 5.30 | 21.40 | 1.70 | 22.20 | 9.00 |
| 3855 | 79.20 | 1.39 | 48.20 | 10.80 | 41.00 | 1.60 | 6.40 | 6.30 |
| 4015 | 92.40 | 1.19 | 79.70 | 12.20 | 54.50 | 1.26 | 20.50 | 10.20 |
| 4032 | 85.40 | 1.48 | 18.40 | 8.30 | 58.30 | 1.42 | 6.30 | 8.80 |
| 4037 | 56.60 | 1.49 | 19.80 | 8.60 | 44.40 | 1.40 | 9.60 | 8.50 |
| 4038 | 87.40 | 1.01 | 70.10 | 14.70 | 50.70 | 1.15 | 12.60 | 8.10 |
| 4040 | 87.30 | 1.42 | 27.10 | 9.40 | 59.60 | 1.43 | 6.30 | 8.60 |
| 4050 | 82.80 | 1.20 | 31.60 | 8.80 | 7.40 | 1.18 | 10.00 | 7.50 |
| 4052 | 40.70 | 1.88 | 18.40 | 6.20 | 35.80 | 1.90 | 11.10 | 5.90 |

| | | | | | | | | |
|---|---|---|---|---|---|---|---|---|
| 4053 | 85.00 | 1.27 | 10.10 | 6.90 | 64.40 | 1.49 | 9.30 | 5.80 |
| 4054 | 89.50 | 1.28 | 79.40 | 7.20 | 84.20 | 1.44 | 84.40 | 8.50 |
| 4055 | 75.00 | 1.29 | 38.10 | 9.40 | 31.70 | 1.37 | 8.90 | 7.10 |
| 4062 | 97.80 | 0.89 | 75.50 | 24.50 | 48.30 | 1.20 | 11.50 | 7.50 |
| 4063 | 99.10 | 0.65 | 95.40 | 48.80 | 71.40 | 1.06 | 13.10 | 8.00 |
| 4071 | 83.10 | 1.29 | 20.10 | 7.20 | 27.10 | 1.37 | 10.60 | 7.20 |
| 4073 | 85.80 | 0.97 | 54.80 | 11.10 | 26.70 | 1.13 | 14.30 | 7.80 |
| 4074 | 37.90 | 1.43 | 13.40 | 6.30 | 14.30 | 1.43 | 7.50 | 8.10 |
| 4076 | 61.60 | 1.27 | 14.90 | 6.40 | 17.00 | 1.35 | 8.90 | 8.20 |
| 4077 | 49.00 | 1.42 | 15.30 | 6.30 | 19.50 | 1.47 | 9.20 | 7.50 |
| 4079 | 70.40 | 1.29 | 13.00 | 6.70 | 18.50 | 1.28 | 9.30 | 7.80 |
| 4080 | 78.00 | 1.13 | 30.50 | 8.30 | 24.00 | 1.23 | 11.50 | 8.30 |
| 4093 | 43.20 | 1.20 | 44.90 | 11.60 | 58.40 | 1.34 | 9.50 | 7.90 |
| 4112 | 79.60 | 1.22 | 15.90 | 6.30 | 56.50 | 1.66 | 7.70 | 4.50 |
| 4114 | 95.40 | 0.69 | 88.40 | 32.30 | 72.50 | 0.95 | 13.80 | 8.30 |
| 4115 | 89.90 | 0.88 | 42.60 | 12.40 | 62.90 | 1.29 | 7.10 | 6.00 |
| 4118 | 48.80 | 1.54 | 13.90 | 6.20 | 38.00 | 1.68 | 6.70 | 5.80 |
| 4121 | 83.40 | 1.06 | 37.20 | 9.50 | 8.60 | 1.08 | 10.20 | 6.80 |
| 4124 | 85.50 | 0.99 | 56.20 | 12.00 | 14.40 | 1.06 | 16.30 | 7.70 |
| 4125 | 84.80 | 0.98 | 54.90 | 11.30 | 14.80 | 1.06 | 12.60 | 7.10 |
| 4128 | 93.30 | 1.04 | 74.70 | 15.90 | 67.50 | 1.18 | 14.00 | 9.50 |
| 4146 | 86.80 | 1.13 | 69.50 | 14.20 | 54.00 | 1.19 | 9.90 | 8.30 |
| 4147 | 82.90 | 1.07 | 59.70 | 13.40 | 47.40 | 1.24 | 8.90 | 7.60 |
| 4148 | 75.80 | 1.31 | 31.90 | 9.40 | 38.00 | 1.35 | 6.90 | 6.90 |
| 5137 | 70.50 | 1.79 | 10.70 | 6.40 | 51.30 | 1.79 | 8.80 | 6.70 |
| 5185 | 94.80 | 1.05 | 86.40 | 13.40 | 68.40 | 1.14 | 13.20 | 6.60 |
| 5415 | 78.90 | 1.69 | 11.80 | 6.20 | 50.10 | 1.79 | 6.90 | 5.10 |
| 5499 | 85.90 | 1.24 | 21.40 | 7.40 | 59.80 | 1.45 | 7.70 | 6.20 |
| 5520 | 18.00 | 1.87 | 10.80 | 6.80 | 18.70 | 1.35 | 6.60 | 9.60 |
| 5600 | 89.20 | 1.50 | 9.20 | 6.30 | 68.00 | 1.59 | 6.10 | 6.40 |
| 5623 | 94.10 | 0.71 | 95.00 | 25.10 | 72.40 | 0.87 | 35.60 | 9.10 |
| 5776 | 45.40 | 1.59 | 16.70 | 9.30 | 50.30 | 1.28 | 6.00 | 11.50 |
| 5777 | 45.60 | 1.68 | 18.10 | 10.00 | 54.70 | 1.32 | 6.50 | 10.50 |
| 5830 | 89.00 | 1.39 | 39.50 | 13.50 | 34.10 | 1.17 | 9.70 | 7.80 |
| 5925 | 94.80 | 1.06 | 52.20 | 23.00 | 11.30 | 0.80 | 27.30 | 11.00 |
| 5995 | 97.40 | 0.81 | 96.40 | 33.20 | 72.90 | 1.12 | 10.20 | 8.30 |
| 6057 | 72.20 | 1.33 | 30.40 | 8.10 | 2.90 | 1.40 | 8.10 | 6.60 |
| 6123 | 65.50 | 1.99 | 42.10 | 19.00 | 72.40 | 0.92 | 90.50 | 10.50 |
| 6124 | 93.20 | 0.99 | 42.20 | 13.60 | 44.30 | 1.28 | 4.90 | 6.50 |
| 6204 | 62.90 | 1.47 | 23.50 | 7.60 | 56.70 | 1.42 | 7.70 | 7.40 |
| 6224 | 94.30 | 0.65 | 90.50 | 36.30 | 71.90 | 0.99 | 19.10 | 9.20 |
| 6239 | 98.50 | 0.80 | 89.50 | 33.40 | 74.80 | 0.99 | 20.40 | 8.90 |
| 6240 | 82.10 | 1.01 | 87.80 | 28.50 | 69.40 | 0.82 | 25.50 | 11.30 |
| 6266 | 64.70 | 0.97 | 68.00 | 17.30 | - | - | - | - |
| 6267 | 76.40 | 1.68 | 19.20 | 8.90 | 67.00 | 1.31 | 5.10 | 8.30 |
| 6270 | 95.80 | 0.94 | 88.10 | 29.30 | 10.30 | 1.04 | 26.30 | 19.00 |
| 6271 | 93.70 | 0.93 | 81.30 | 15.60 | 11.70 | 1.20 | 13.30 | 9.00 |
| 6272 | 97.00 | 0.71 | 92.50 | 24.10 | 88.90 | 0.70 | 73.10 | 19.60 |
| 6310 | 86.70 | 1.43 | 62.20 | 15.90 | 39.80 | 1.50 | 7.80 | 12.80 |
| 6311 | 78.50 | 1.43 | 28.00 | 7.50 | 4.90 | 1.20 | 8.60 | 7.60 |
| 6324 | 84.30 | 1.06 | 79.50 | 15.70 | 69.40 | 1.13 | 9.10 | 8.20 |
| 6337 | 86.80 | 1.08 | 41.30 | 11.60 | 59.70 | 1.37 | 7.70 | 7.20 |
| 6338 | 88.00 | 1.12 | 48.00 | 12.90 | 66.50 | 1.22 | 8.10 | 8.10 |
| 6344 | 79.50 | 1.12 | 19.80 | 10.70 | 75.10 | 1.12 | 6.30 | 8.70 |
| 6345 | 80.50 | 1.14 | 21.00 | 11.30 | 79.60 | 1.32 | 5.20 | 9.00 |
| 6346 | 84.80 | 1.08 | 55.20 | 13.80 | 72.80 | 1.11 | 18.10 | 10.00 |
| 6349 | 97.50 | 0.76 | 94.40 | 37.80 | 45.90 | 1.54 | 6.30 | 6.90 |
| 6350 | 93.00 | 0.82 | 81.90 | 22.00 | 41.90 | 1.70 | 6.60 | 5.80 |
| 6351 | 94.50 | 0.78 | 89.30 | 32.50 | 43.40 | 1.01 | 11.00 | 9.50 |
| 6352 | 98.00 | 0.85 | 89.10 | 26.20 | 26.70 | 1.19 | 8.80 | 6.70 |
| 6353 | 97.80 | 0.81 | 92.90 | 27.80 | 12.80 | 1.24 | 6.40 | 6.40 |
| 6354 | 94.70 | 0.90 | 70.80 | 17.30 | 30.10 | 1.80 | 3.10 | 6.50 |
| 6394 | 96.60 | 0.94 | 87.70 | 24.10 | 67.90 | 1.42 | 12.70 | 6.80 |
| 6398 | 97.40 | 0.77 | 95.00 | 29.70 | 51.20 | 1.33 | 12.20 | 6.70 |
| 6404 | 90.70 | 1.09 | 72.30 | 13.50 | 58.10 | 1.34 | 9.80 | 7.20 |

| | | | | | | | | |
|---|---|---|---|---|---|---|---|---|
| 6408 | 97.10 | 0.83 | 93.50 | 28.50 | 79.30 | 0.83 | 24.30 | 10.40 |
| 6413 | 58.40 | 1.38 | 35.50 | 8.50 | 21.50 | 1.34 | 8.60 | 7.50 |
| 6414 | 47.50 | 1.42 | 53.20 | 10.50 | 22.90 | 1.30 | 10.70 | 8.30 |
| 6415 | 38.70 | 1.75 | 50.10 | 9.80 | 20.10 | 1.71 | 13.10 | 7.60 |
| 6416 | 39.20 | 1.40 | 46.90 | 10.10 | 20.60 | 1.43 | 13.80 | 8.30 |
| 6417 | 32.40 | 1.38 | 41.20 | 8.70 | 16.60 | 1.47 | 11.70 | 7.10 |
| 6418 | 48.70 | 1.49 | 29.40 | 8.10 | 20.80 | 1.42 | 8.30 | 8.10 |
| 6425 | 50.90 | 1.76 | 9.40 | 5.70 | 43.70 | 1.84 | 4.10 | 4.30 |
| 6433 | 76.60 | 0.94 | 19.60 | 6.30 | 51.60 | 1.50 | 6.30 | 5.70 |
| 6435 | 89.90 | 1.00 | 49.50 | 9.30 | 65.70 | 1.34 | 9.40 | 7.00 |
| 6455 | 71.00 | 1.35 | 60.10 | 11.60 | 53.90 | 1.11 | 8.20 | 11.70 |
| 6475 | 56.00 | 1.52 | 11.90 | 7.80 | 45.40 | 1.47 | 5.60 | 8.50 |
| 6480 | 46.60 | 1.35 | 60.60 | 13.50 | 25.80 | 1.32 | 8.70 | 8.10 |
| 6481 | 71.70 | 1.76 | 14.50 | 6.80 | 26.90 | 1.65 | 5.10 | 8.00 |
| 6483 | 73.20 | 1.35 | 22.30 | 7.50 | 9.20 | 1.26 | 14.80 | 6.80 |
| 6486 | 79.30 | 1.20 | 28.90 | 8.20 | 12.20 | 1.14 | 16.40 | 7.30 |
| 6487 | 94.70 | 0.99 | 66.90 | 14.20 | 60.00 | 1.33 | 6.50 | 6.40 |
| 6488 | 93.30 | 1.02 | 74.60 | 14.00 | 57.30 | 1.19 | 7.90 | 7.40 |
| 6526 | 71.70 | 1.22 | 40.50 | 9.00 | 38.00 | 1.37 | 9.10 | 7.60 |
| 6534 | 81.60 | 1.22 | 46.00 | 10.70 | 49.80 | 1.33 | 7.30 | 7.30 |
| 6551 | 95.40 | 1.07 | 86.80 | 22.20 | 73.90 | 1.13 | 19.80 | 9.60 |
| 6555 | 100.00 | 0.56 | 97.90 | 95.00 | 73.20 | 0.94 | 17.30 | 10.70 |
| 6559 | 79.30 | 1.30 | 41.90 | 10.00 | 14.70 | 1.29 | 12.40 | 7.50 |
| 6561 | 50.40 | 1.16 | 50.50 | 11.90 | 24.50 | 1.29 | 10.50 | 7.50 |
| 6562 | 50.70 | 1.20 | 49.20 | 11.30 | 23.80 | 1.31 | 9.50 | 7.60 |
| 6563 | 42.50 | 1.21 | 58.70 | 12.30 | 23.10 | 1.17 | 10.90 | 8.80 |
| 6564 | 31.50 | 1.44 | 39.00 | 9.10 | 13.10 | 1.47 | 10.80 | 7.90 |
| 6565 | 29.50 | 1.24 | 42.30 | 11.60 | 19.20 | 1.41 | 10.90 | 7.90 |
| 6566 | 33.60 | 1.28 | 46.80 | 10.80 | 14.80 | 1.38 | 10.00 | 7.50 |
| 6567 | 28.80 | 1.36 | 40.20 | 10.00 | 15.30 | 1.31 | 9.00 | 8.70 |
| 6568 | 20.40 | 1.34 | 42.40 | 9.90 | 12.20 | 1.45 | 8.00 | 8.40 |
| 6569 | 24.50 | 1.46 | 26.50 | 8.80 | 11.90 | 1.56 | 8.10 | 6.90 |
| 6570 | 22.30 | 1.67 | 25.20 | 7.60 | 10.70 | 1.61 | 5.70 | 7.00 |
| 6573 | 37.20 | 1.22 | 48.10 | 11.80 | 17.80 | 1.32 | 11.30 | 8.30 |
| 6580 | 71.40 | 1.73 | 16.50 | 8.30 | 60.10 | 1.67 | 5.80 | 7.40 |
| 6583 | 80.60 | 0.85 | 80.10 | 17.50 | 45.00 | 0.86 | 15.60 | 10.40 |
| 6584 | 73.00 | 1.32 | 27.80 | 7.40 | 3.20 | 1.17 | 13.80 | 7.80 |
| 6615 | 90.50 | 0.89 | 73.50 | 17.00 | 38.10 | 0.96 | 17.00 | 8.80 |
| 6617 | 79.80 | 0.90 | 79.30 | 15.70 | 50.70 | 0.93 | 15.60 | 10.40 |
| 6629 | 59.10 | 1.13 | 16.50 | 7.20 | 22.90 | 1.71 | 6.80 | 5.00 |
| 6630 | 92.90 | 0.83 | 90.20 | 21.00 | 66.10 | 1.05 | 38.70 | 9.60 |
| 6631 | 87.30 | 0.86 | 83.10 | 20.60 | 59.50 | 1.07 | 12.90 | 8.00 |
| 6632 | 82.70 | 1.14 | 72.20 | 12.40 | 53.40 | 1.04 | 14.30 | 9.10 |
| 6633 | 90.70 | 0.88 | 87.10 | 18.80 | 67.20 | 0.97 | 17.70 | 9.80 |
| 6634 | 89.10 | 0.96 | 72.60 | 18.40 | 66.70 | 0.91 | 12.70 | 10.30 |
| 6635 | 88.50 | 0.84 | 74.90 | 19.10 | 62.70 | 0.89 | 36.30 | 10.70 |
| 6637 | 92.70 | 0.80 | 36.60 | 8.70 | 72.60 | 1.41 | 6.40 | 7.20 |
| 6640 | 69.90 | 1.63 | 10.00 | 8.40 | 55.70 | 1.65 | 5.20 | 6.90 |
| 6656 | 92.50 | 0.98 | 83.70 | 14.80 | 62.70 | 1.05 | 15.00 | 8.20 |
| 6667 | 86.70 | 0.76 | 87.20 | 22.10 | 43.70 | 0.82 | 13.40 | 9.70 |
| 6668 | 42.80 | 1.77 | 8.80 | 7.10 | 40.80 | 1.50 | 6.70 | 8.70 |
| 6675 | 73.90 | 0.66 | 87.40 | 38.80 | 56.10 | 0.83 | 20.40 | 13.90 |
| 6676 | 76.60 | 0.75 | 88.20 | 38.00 | 54.20 | 0.93 | 15.60 | 10.30 |
| 6677 | 68.20 | 1.38 | 51.20 | 11.60 | 47.70 | 1.49 | 11.50 | 7.80 |
| 6679 | 66.30 | 1.44 | 34.60 | 8.00 | 48.90 | 1.48 | 7.80 | 7.80 |
| 6685 | 70.60 | 1.41 | 17.00 | 6.20 | 21.30 | 1.79 | 8.20 | 5.50 |
| 6686 | 97.90 | 0.60 | 95.50 | 50.60 | 73.60 | 1.04 | 15.80 | 9.20 |
| 6687 | 96.10 | 0.88 | 64.90 | 16.20 | 63.70 | 1.42 | 9.60 | 7.20 |
| 6688 | 66.80 | 1.16 | 17.70 | 6.30 | 41.70 | 1.46 | 5.00 | 5.80 |
| 6694 | 52.70 | 1.68 | 11.30 | 6.70 | 44.80 | 1.70 | 8.10 | 6.90 |
| 6698 | 85.20 | 1.64 | 65.20 | 11.20 | 63.10 | 1.50 | 8.00 | 9.80 |
| 6699 | 77.20 | 1.28 | 49.30 | 9.90 | 36.80 | 1.10 | 12.10 | 8.50 |
| 6703 | 83.60 | 0.97 | 68.80 | 14.00 | 53.20 | 1.28 | 14.60 | 7.70 |
| 6704 | 69.40 | 1.70 | 32.60 | 7.10 | 41.10 | 1.73 | 8.50 | 6.70 |
| 6705 | 74.30 | 1.18 | 50.50 | 9.70 | 14.80 | 1.26 | 8.40 | 7.00 |

| | | | | | | | | |
|---|---|---|---|---|---|---|---|---|
| 6706 | 59.20 | 1.54 | 27.70 | 6.60 | 11.30 | 1.33 | 13.20 | 7.20 |
| 6707 | 59.10 | 1.59 | 23.30 | 7.10 | 39.50 | 1.62 | 7.50 | 5.80 |
| 6709 | 72.60 | 1.16 | 43.20 | 9.80 | 8.00 | 1.08 | 12.10 | 7.60 |
| 6710 | 71.50 | 1.11 | 33.60 | 10.80 | 29.30 | 1.15 | 11.00 | 9.10 |
| 6711 | 87.70 | 1.02 | 73.00 | 14.40 | 11.70 | 0.98 | 14.50 | 8.90 |
| 6721 | 47.90 | 1.16 | 61.60 | 13.20 | 25.10 | 1.19 | 10.90 | 7.90 |
| 6724 | 71.20 | 1.29 | 33.60 | 10.70 | 50.60 | 1.29 | 7.40 | 9.50 |
| 6732 | 79.70 | 1.12 | 46.60 | 9.50 | 51.20 | 1.29 | 12.60 | 6.90 |
| 6741 | 85.00 | 0.79 | 85.00 | 19.80 | 53.10 | 0.72 | 30.90 | 10.60 |
| 6742 | 86.40 | 0.93 | 76.50 | 15.30 | 45.70 | 0.89 | 13.70 | 9.60 |
| 6743 | 90.20 | 0.96 | 59.90 | 11.50 | 58.30 | 1.30 | 12.90 | 6.60 |
| 6744 | 85.00 | 1.05 | 43.70 | 10.50 | 55.40 | 1.22 | 8.90 | 8.30 |
| 6747 | 68.40 | 0.95 | 73.80 | 17.70 | 41.00 | 1.29 | 12.80 | 8.20 |
| 6751 | 98.90 | 0.66 | 91.10 | 44.20 | 87.10 | 0.87 | 43.30 | 13.00 |
| 6752 | 98.50 | 0.65 | 89.50 | 35.40 | 80.20 | 0.93 | 47.30 | 11.70 |
| 6757 | 97.20 | 0.73 | 90.90 | 23.20 | 69.90 | 1.37 | 8.80 | 6.50 |
| 6770 | 87.90 | 1.25 | 51.80 | 12.10 | 68.50 | 1.21 | 8.10 | 9.10 |
| 6771 | 92.80 | 1.12 | 78.80 | 16.40 | 56.20 | 1.17 | 13.60 | 7.90 |
| 6772 | 89.90 | 1.09 | 72.60 | 16.80 | 59.90 | 1.13 | 10.20 | 8.90 |
| 6773 | 68.00 | 1.19 | 58.10 | 11.80 | 42.40 | 1.13 | 9.10 | 9.40 |
| 6774 | 86.50 | 0.94 | 85.60 | 19.40 | 61.70 | 1.02 | 18.70 | 9.00 |
| 6777 | 89.90 | 0.94 | 86.20 | 21.80 | 64.20 | 0.85 | 25.00 | 9.40 |
| 6778 | 94.70 | 0.82 | 81.80 | 19.50 | 48.40 | 1.02 | 21.80 | 8.80 |
| 6780 | 87.80 | 0.95 | 62.30 | 13.70 | 38.50 | 1.13 | 19.60 | 8.90 |
| 6784 | 94.00 | 0.83 | 81.80 | 20.20 | 47.60 | 0.98 | 18.20 | 8.90 |
| 6788 | 89.90 | 0.99 | 58.20 | 12.10 | 37.20 | 1.15 | 15.10 | 8.20 |
| 6789 | 94.20 | 0.81 | 84.10 | 20.20 | 46.60 | 1.02 | 15.30 | 9.10 |
| 6790 | 92.40 | 0.93 | 58.40 | 12.90 | 32.30 | 1.02 | 12.10 | 8.40 |
| 7018 | 88.00 | 1.10 | 63.20 | 13.50 | 60.90 | 1.13 | 10.00 | 8.80 |
| 7019 | 88.20 | 1.14 | 74.50 | 16.20 | 62.60 | 1.04 | 17.80 | 8.30 |
| 7030 | 81.30 | 1.35 | 27.20 | 8.20 | 2.60 | 1.61 | 9.30 | 5.40 |
| 7035 | 67.20 | 1.53 | 10.10 | 5.70 | 3.20 | 2.05 | 5.00 | 4.00 |
| 7036 | 91.60 | 1.17 | 72.90 | 18.60 | 63.70 | 1.13 | 14.60 | 11.00 |
| 7040 | 92.70 | 1.42 | 26.60 | 9.70 | 59.00 | 1.28 | 9.90 | 7.20 |
| 7048 | 76.30 | 1.07 | 61.60 | 12.50 | 36.90 | 1.03 | 27.50 | 8.90 |
| 7049 | 71.80 | 1.30 | 21.80 | 7.10 | 26.10 | 1.37 | 11.50 | 6.60 |
| 7050 | 83.80 | 0.88 | 69.00 | 15.90 | 44.60 | 1.07 | 25.20 | 9.60 |
| 7051 | 81.30 | 1.03 | 83.00 | 23.60 | 47.30 | 1.16 | 28.90 | 9.30 |
| 7052 | 65.30 | 1.02 | 60.30 | 11.40 | 30.40 | 1.29 | 16.60 | 6.50 |
| 7063 | 91.00 | 0.85 | 77.50 | 17.20 | 71.20 | 1.20 | 12.60 | 7.60 |
| 7073 | 75.70 | 1.21 | 61.90 | 13.90 | 55.30 | 1.22 | 9.60 | 11.20 |
| 8000 | 66.10 | 0.95 | 48.80 | 9.50 | 5.20 | 1.14 | 14.40 | 9.00 |
| 8001 | 72.10 | 0.84 | 77.50 | 15.50 | 1.60 | 0.83 | 14.30 | 21.00 |
| 8002 | 64.50 | 1.65 | 18.60 | 6.40 | 0.30 | 2.01 | 16.70 | 4.00 |
| 8003 | 80.00 | 0.90 | 53.20 | 12.00 | 12.30 | 1.02 | 12.50 | 9.70 |
| 8004 | 82.40 | 1.07 | 55.90 | 11.60 | 4.40 | 0.88 | 16.10 | 20.10 |
| 8011 | 84.00 | 1.46 | 12.80 | 6.40 | 37.80 | 1.40 | 7.00 | 7.30 |
| 8012 | 61.40 | 1.47 | 29.90 | 7.90 | 28.50 | 1.22 | 7.00 | 9.10 |
| 8013 | 77.70 | 1.70 | 7.60 | 5.60 | 19.80 | 1.44 | 7.70 | 7.00 |
| 8014 | 91.00 | 1.18 | 45.40 | 12.40 | 39.60 | 1.17 | 8.20 | 8.30 |
| 8015 | 97.60 | 0.62 | 94.30 | 46.80 | 79.20 | 1.10 | 10.10 | 12.00 |
| 8064 | 73.10 | 1.10 | 56.00 | 11.30 | 40.60 | 1.38 | 7.50 | 6.50 |
| 8069 | 92.90 | 1.14 | 38.30 | 10.30 | 66.70 | 1.51 | 5.30 | 6.10 |
| 8072 | 97.30 | 0.72 | 94.90 | 35.80 | 84.60 | 1.07 | 20.00 | 9.00 |
| 8094 | 83.50 | 1.14 | 18.30 | 8.10 | 37.10 | 1.54 | 4.40 | 5.90 |
| 8095 | 92.60 | 1.00 | 51.70 | 12.20 | 53.60 | 1.27 | 7.80 | 8.70 |
| 8099 | 77.70 | 0.87 | 31.70 | 9.70 | 52.10 | 1.45 | 3.00 | 6.00 |
| 8100 | 82.20 | 0.92 | 32.40 | 11.10 | 51.20 | 1.40 | 8.60 | 6.40 |
| 8107 | 80.10 | 0.92 | 81.00 | 19.00 | 34.20 | 1.02 | 13.10 | 8.70 |
| 8116 | 84.90 | 1.11 | 18.80 | 7.50 | 25.20 | 1.37 | 5.50 | 6.60 |
| 8117 | 90.80 | 0.98 | 82.70 | 17.00 | 71.60 | 0.95 | 46.00 | 10.10 |
| 8118 | 90.10 | 1.06 | 78.00 | 17.80 | 68.80 | 0.87 | 13.70 | 9.30 |
| 8119 | 81.20 | 1.11 | 78.40 | 18.80 | 58.40 | 0.85 | 20.90 | 11.00 |
| 8124 | 85.30 | 1.20 | 40.20 | 9.80 | 32.50 | 1.38 | 11.40 | 7.00 |
| 8136 | 55.00 | 1.03 | 39.50 | 10.70 | 38.10 | 1.31 | 8.80 | 7.30 |

| | | | | | | | | |
|---|---|---|---|---|---|---|---|---|
| 8137 | 53.90 | 1.02 | 45.00 | 11.10 | 36.70 | 1.26 | 9.80 | 7.50 |
| 8138 | 59.80 | 0.98 | 44.10 | 11.60 | 30.80 | 1.32 | 8.90 | 7.90 |
| 8150 | 90.40 | 1.16 | 17.20 | 10.40 | 42.60 | 1.36 | 6.30 | 7.90 |
| 8162 | 92.90 | 0.92 | 79.10 | 16.20 | 68.30 | 1.22 | 8.00 | 6.40 |
| 8163 | 94.50 | 0.93 | 72.00 | 15.70 | 63.90 | 1.26 | 9.80 | 5.80 |
| 8177 | 37.60 | 1.31 | 62.40 | 11.00 | 32.90 | 1.53 | 5.20 | 7.60 |
| 8178 | 31.60 | 1.19 | 73.80 | 12.60 | 28.00 | 1.34 | 5.30 | 7.00 |
| 8179 | 95.30 | 1.06 | 73.70 | 22.50 | 40.00 | 1.47 | 12.10 | 6.20 |
| 8183 | 83.00 | 1.27 | 31.70 | 7.00 | 54.20 | 1.29 | 11.20 | 8.40 |
| 8184 | 93.50 | 0.64 | 92.00 | 37.60 | 73.30 | 1.00 | 11.40 | 9.00 |
| 8188 | 79.40 | 1.32 | 30.60 | 8.20 | 28.10 | 1.46 | 5.90 | 5.90 |
| 8189 | 95.70 | 0.58 | 96.70 | 34.50 | 87.40 | 0.83 | 63.20 | 16.10 |
| 8191 | 84.50 | 0.98 | 40.70 | 12.80 | 71.30 | 1.04 | 15.00 | 7.50 |
| 8192 | 63.20 | 1.34 | 15.10 | 6.80 | 59.20 | 1.26 | 8.90 | 7.30 |
| 8193 | 73.60 | 1.21 | 13.00 | 7.90 | 69.50 | 1.16 | 10.50 | 8.60 |
| 8194 | 96.90 | 0.69 | 93.00 | 47.50 | 78.60 | 0.79 | 36.80 | 9.60 |
| 8200 | 75.20 | 1.11 | 48.10 | 10.10 | 71.70 | 1.29 | 7.80 | 8.90 |
| 8215 | 53.60 | 1.34 | 34.50 | 8.40 | 55.40 | 1.31 | 4.50 | 8.50 |
| 8240 | 69.50 | 1.37 | 20.90 | 5.90 | 35.60 | 1.51 | 6.10 | 5.60 |
| 8242 | 7.90 | 2.02 | 10.20 | 6.40 | 29.30 | 1.93 | 7.10 | 4.80 |
| 8253 | 91.00 | 0.86 | 80.80 | 16.90 | 5.90 | 1.54 | 4.30 | 11.50 |
| 8266 | 90.20 | 0.90 | 77.20 | 27.90 | 22.50 | 1.34 | 5.40 | 8.00 |
| 8284 | 33.00 | 1.35 | 11.30 | 4.90 | 50.20 | 1.65 | 5.00 | 5.80 |
| 8289 | 82.40 | 1.57 | 23.50 | 7.10 | 50.70 | 1.40 | 7.90 | 7.00 |
| 8298 | 93.40 | 0.93 | 63.70 | 12.60 | 64.50 | 1.72 | 8.30 | 5.80 |
| 8315 | 84.40 | 1.31 | 59.30 | 11.20 | 53.60 | 1.21 | 14.10 | 9.80 |
| 8320 | 96.10 | 0.91 | 82.40 | 19.00 | - | - | - | - |
| 8321 | 94.30 | 1.06 | 75.90 | 15.80 | 40.40 | 1.38 | 6.10 | 6.10 |
| 8322 | 88.00 | 1.14 | 23.70 | 7.50 | 21.80 | 1.22 | 4.80 | 7.20 |
| 8323 | 75.20 | 1.39 | 10.30 | 7.30 | 33.50 | 1.56 | 7.40 | 6.10 |
| 8331 | 94.40 | 0.87 | 76.70 | 20.20 | 68.60 | 1.04 | 12.00 | 8.60 |
| 8334 | 64.80 | 1.97 | 9.10 | 5.90 | 47.80 | 1.93 | 7.60 | 6.00 |
| 8342 | 76.30 | 1.32 | 51.20 | 10.10 | 46.00 | 1.20 | 10.60 | 8.20 |
| 8354 | 96.10 | 0.75 | 95.50 | 40.50 | 79.70 | 0.85 | 24.40 | 9.20 |
| 8361 | 93.70 | 0.78 | 85.80 | 22.50 | 58.00 | 0.83 | 23.70 | 10.90 |
| 8368 | 75.20 | 1.22 | 19.40 | 7.50 | 23.30 | 1.30 | 8.70 | 7.20 |
| 8372 | 78.10 | 1.42 | 40.40 | 9.20 | 25.20 | 1.27 | 6.00 | 8.80 |
| 8373 | 77.70 | 1.31 | 51.40 | 10.80 | 44.80 | 1.31 | 8.20 | 8.10 |
| 8375 | 66.50 | 1.55 | 29.00 | 7.80 | 13.30 | 1.44 | 4.90 | 8.50 |
| 8376 | 53.10 | 1.54 | 25.70 | 8.10 | 23.30 | 1.45 | 5.70 | 8.20 |
| 8377 | 54.40 | 1.49 | 29.70 | 8.10 | 13.50 | 1.45 | 4.80 | 8.40 |
| 8378 | 60.00 | 1.42 | 39.00 | 8.90 | 12.90 | 1.40 | 5.20 | 9.20 |
| 8379 | 63.40 | 1.44 | 35.60 | 8.70 | 41.00 | 1.29 | 5.00 | 9.30 |
| 8380 | 67.60 | 1.42 | 36.50 | 8.80 | 6.50 | 1.28 | 4.90 | 9.20 |
| 8381 | 61.90 | 1.50 | 30.60 | 8.10 | 39.80 | 1.47 | 5.70 | 8.00 |
| 8382 | 71.90 | 1.39 | 41.90 | 9.20 | 33.40 | 1.27 | 5.70 | 8.90 |
| 8387 | 53.70 | 1.51 | 28.50 | 8.30 | 42.30 | 1.43 | 4.80 | 7.90 |
| 8390 | 63.40 | 1.56 | 19.80 | 7.80 | 46.10 | 1.49 | 4.60 | 8.30 |
| 8391 | 66.60 | 1.42 | 37.00 | 8.70 | 34.70 | 1.46 | 7.40 | 7.40 |
| 8395 | 73.00 | 1.44 | 31.60 | 8.60 | 45.20 | 1.41 | 4.30 | 8.40 |
| 8397 | 95.80 | 0.70 | 84.30 | 35.00 | 66.30 | 1.08 | 11.90 | 7.60 |
| 8398 | 94.40 | 0.87 | 80.70 | 18.80 | 64.80 | 1.43 | 5.70 | 6.40 |
| 8399 | 90.20 | 0.73 | 86.50 | 27.50 | 68.20 | 1.12 | 10.00 | 8.50 |
| 8405 | 86.60 | 0.99 | 29.60 | 8.90 | 82.30 | 1.21 | 30.40 | 7.50 |
| 8409 | 91.10 | 1.36 | 35.90 | 12.70 | 60.70 | 1.40 | 10.20 | 10.00 |
| 8410 | 90.80 | 1.01 | 80.20 | 15.20 | 67.60 | 1.07 | 12.60 | 8.50 |
| 8414 | 24.80 | 1.63 | 12.80 | 6.30 | 61.30 | 1.30 | 9.20 | 7.30 |
| 8435 | 91.00 | 0.99 | 67.30 | 14.40 | 65.70 | 1.17 | 11.10 | 7.60 |
| 8454 | 82.30 | 1.12 | 54.40 | 9.20 | 65.40 | 1.31 | 9.50 | 7.90 |
| 8461 | 82.20 | 1.45 | 55.90 | 11.10 | 65.70 | 1.48 | 7.10 | 7.70 |
| 8469 | 71.60 | 1.73 | 9.60 | 8.30 | 47.70 | 1.73 | 9.20 | 6.80 |
| 8477 | 84.00 | 1.09 | 66.50 | 13.60 | 31.30 | 1.24 | 15.40 | 7.80 |
| 8478 | 91.50 | 0.93 | 83.30 | 20.70 | 56.70 | 1.12 | 17.50 | 8.50 |
| 8479 | 47.70 | 1.64 | 11.70 | 7.30 | 43.30 | 1.60 | 6.50 | 7.20 |
| 8481 | 87.90 | 1.09 | 35.20 | 8.70 | 44.40 | 1.30 | 11.20 | 7.50 |

| | | | | | | | | |
|---|---|---|---|---|---|---|---|---|
| 8482 | 75.90 | 1.37 | 20.90 | 9.30 | 65.90 | 1.41 | 6.00 | 9.70 |
| 8505 | 53.70 | 1.81 | 9.00 | 5.80 | 2.50 | 1.42 | 8.60 | 7.20 |
| 8506 | 86.80 | 1.14 | 34.10 | 9.30 | 7.10 | 1.16 | 10.80 | 7.40 |
| 8511 | 90.90 | 1.21 | 73.10 | 13.30 | 67.60 | 1.00 | 25.20 | 9.30 |
| 8512 | 95.00 | 1.10 | 78.90 | 17.70 | 50.60 | 1.00 | 21.20 | 8.20 |
| 8515 | 94.00 | 0.96 | 63.40 | 16.00 | 76.50 | 1.19 | 5.40 | 8.40 |
| 8516 | 81.10 | 1.42 | 46.80 | 9.50 | 64.00 | 1.48 | 6.00 | 8.40 |
| 8517 | 78.60 | 1.34 | 13.60 | 8.10 | 67.70 | 1.63 | 6.70 | 6.80 |
| 8540 | 54.70 | 1.38 | 34.50 | 9.40 | 38.00 | 1.50 | 8.30 | 7.10 |
| 8559 | 86.00 | 1.33 | 61.10 | 11.50 | 62.40 | 1.23 | 12.00 | 7.80 |
| 8560 | 88.40 | 1.43 | 50.30 | 9.20 | 67.80 | 1.17 | 14.10 | 8.90 |
| 8574 | 96.30 | 0.54 | 96.20 | 41.50 | 72.10 | 0.90 | 38.30 | 10.10 |
| 8576 | 94.80 | 0.87 | 80.10 | 18.30 | 50.30 | 0.95 | 14.20 | 9.00 |
| 8579 | 7.70 | 2.05 | 10.20 | 6.40 | 45.10 | 1.83 | 7.90 | 6.00 |
| 8598 | 88.80 | 0.75 | 88.30 | 23.80 | 72.20 | 1.09 | 29.60 | 8.60 |
| 8605 | 92.90 | 0.79 | 89.20 | 29.50 | 72.20 | 1.18 | 15.00 | 8.30 |
| 8608 | 84.40 | 0.92 | 64.40 | 12.50 | 49.30 | 1.42 | 6.30 | 5.30 |
| 8623 | 90.60 | 1.16 | 38.80 | 11.80 | 67.90 | 1.34 | 8.00 | 7.90 |
| 8632 | 64.10 | 1.36 | 56.50 | 13.10 | 57.40 | 1.20 | 12.20 | 14.80 |
| 8633 | 47.30 | 1.43 | 14.20 | 15.20 | 63.30 | 1.29 | 21.20 | 11.10 |
| 8637 | 93.30 | 1.32 | 36.10 | 18.00 | 80.00 | 1.37 | 3.20 | 15.40 |
| 8641 | 77.90 | 1.06 | 47.90 | 10.00 | 19.20 | 1.19 | 11.40 | 7.60 |
| 8642 | 90.20 | 1.34 | 59.50 | 10.60 | 76.30 | 1.28 | 10.00 | 9.40 |
| 8643 | 82.70 | 1.23 | 26.50 | 7.80 | 48.30 | 1.40 | 6.20 | 6.10 |
| 8644 | 77.80 | 1.17 | 14.30 | 6.40 | 65.30 | 1.47 | 4.90 | 6.00 |
| 8645 | 63.20 | 1.27 | 20.90 | 7.10 | 12.70 | 1.22 | 10.90 | 7.30 |
| 8650 | 58.60 | 1.41 | 17.40 | 9.90 | 62.50 | 1.26 | 10.60 | 9.00 |
| 8651 | 61.30 | 1.54 | 13.50 | 7.80 | 70.30 | 1.44 | 10.20 | 7.70 |
| 8652 | 52.80 | 1.56 | 11.90 | 8.20 | 57.40 | 1.44 | 6.20 | 8.20 |
| 8653 | 73.40 | 1.57 | 6.60 | 6.50 | 58.60 | 1.70 | 8.80 | 6.00 |
| 8658 | 88.10 | 1.34 | 41.10 | 8.70 | 68.20 | 1.49 | 8.60 | 6.60 |
| 8672 | 56.90 | 1.88 | 10.20 | 6.40 | 47.10 | 1.86 | 7.10 | 7.20 |
| 8697 | 83.00 | 1.24 | 52.60 | 10.40 | 59.50 | 1.37 | 7.80 | 7.60 |
| 8702 | 70.80 | 1.20 | 54.50 | 10.20 | 48.20 | 1.12 | 11.40 | 8.50 |
| 8708 | 88.40 | 0.86 | 78.10 | 16.30 | 29.10 | 0.91 | 16.80 | 8.10 |
| 8712 | 90.90 | 1.09 | 76.40 | 18.30 | 64.90 | 1.22 | 10.20 | 7.10 |
| 8713 | 79.80 | 1.05 | 47.20 | 9.80 | 45.10 | 1.17 | 7.00 | 8.30 |
| 8717 | 88.20 | 1.04 | 46.50 | 9.00 | 45.20 | 1.17 | 7.30 | 8.10 |
| 8732 | 86.70 | 1.06 | 74.60 | 14.10 | 49.80 | 1.20 | 12.90 | 8.30 |
| 8744 | 31.00 | 1.93 | 11.30 | 6.50 | 14.40 | 1.98 | 7.60 | 6.30 |
| 8745 | 22.00 | 2.07 | 10.30 | 5.70 | 12.60 | 1.98 | 8.20 | 6.70 |
| 8746 | 91.50 | 1.17 | 61.30 | 13.60 | 13.00 | 1.08 | 9.30 | 10.70 |
| 8751 | 73.60 | 1.78 | 9.10 | 7.10 | 63.10 | 1.67 | 6.20 | 8.20 |
| 8755 | 92.20 | 1.12 | 62.80 | 14.30 | 45.40 | 1.56 | 8.40 | 6.00 |
| 8756 | 92.20 | 1.02 | 50.90 | 16.30 | 52.90 | 1.45 | 5.00 | 6.60 |
| 8757 | 90.70 | 1.09 | 56.00 | 14.80 | 46.10 | 1.78 | 5.20 | 5.50 |
| 8764 | 92.60 | 1.05 | 78.60 | 14.50 | 66.60 | 0.97 | 42.10 | 10.80 |
| 8767 | 95.70 | 0.85 | 74.40 | 31.20 | 42.30 | 1.45 | 11.60 | 11.50 |
| 8771 | 66.90 | 1.04 | 59.90 | 13.10 | 38.40 | 1.32 | 7.80 | 7.30 |
| 8774 | 48.90 | 1.33 | 14.10 | 7.80 | 30.50 | 1.47 | 6.60 | 7.30 |
| 8775 | 67.20 | 1.13 | 43.50 | 10.20 | 37.40 | 1.40 | 5.70 | 7.50 |
| 8776 | 54.50 | 1.30 | 18.70 | 7.80 | 30.90 | 1.46 | 6.40 | 7.40 |
| 8780 | 70.80 | 1.68 | 11.50 | 7.90 | 77.00 | 1.67 | 9.60 | 9.40 |
| 8782 | 90.50 | 1.41 | 66.50 | 15.90 | 66.70 | 1.38 | 12.30 | 8.10 |
| 8783 | 69.20 | 1.36 | 34.00 | 7.70 | 38.50 | 1.55 | 6.80 | 5.80 |
| 8784 | 84.70 | 1.06 | 69.50 | 12.80 | 54.10 | 1.24 | 6.00 | 7.90 |
| 8790 | 60.40 | 1.43 | 32.90 | 6.70 | 32.90 | 1.63 | 9.50 | 5.80 |
| 8791 | 79.10 | 1.12 | 46.40 | 9.40 | 47.60 | 1.36 | 7.50 | 6.60 |
| 8794 | 81.60 | 1.49 | 8.00 | 7.30 | 64.30 | 1.53 | 6.20 | 6.60 |
| 8795 | 80.60 | 1.04 | 30.40 | 10.20 | 66.20 | 1.50 | 27.20 | 7.10 |
| 8820 | 73.20 | 1.48 | 18.30 | 6.90 | 53.80 | 1.54 | 6.70 | 6.70 |
| 8823 | 79.40 | 1.32 | 36.20 | 9.60 | 46.90 | 1.37 | 6.50 | 9.40 |
| 8827 | 59.60 | 1.13 | 20.40 | 7.90 | 28.10 | 1.39 | 10.60 | 7.00 |
| 8847 | 92.70 | 1.29 | 63.90 | 12.20 | 73.80 | 1.22 | 13.50 | 8.30 |
| 8848 | 87.70 | 1.45 | 49.40 | 12.60 | - | - | - | - |

| | | | | | | | | |
|---|---|---|---|---|---|---|---|---|
| 8850 | 89.70 | 1.23 | 27.70 | 11.30 | 75.20 | 1.39 | 11.50 | 9.50 |
| 8852 | 86.10 | 1.38 | 26.90 | 11.80 | 69.50 | 1.47 | 11.40 | 11.10 |
| 8855 | 91.40 | 1.15 | 30.10 | 17.60 | 79.20 | 1.12 | 15.50 | 16.40 |
| 8860 | 77.80 | 1.17 | 39.60 | 13.40 | 39.40 | 1.16 | 11.50 | 8.70 |
| 8881 | 91.20 | 1.01 | 78.00 | 23.00 | 68.90 | 1.03 | 12.10 | 8.50 |
| 8882 | 94.30 | 1.04 | 78.30 | 17.80 | 68.50 | 1.02 | 21.90 | 8.60 |
| 8883 | 92.30 | 1.01 | 85.00 | 19.00 | 66.70 | 1.14 | 28.10 | 7.60 |
| 9400 | 86.50 | 1.12 | 60.60 | 13.00 | 57.50 | 1.20 | 7.10 | 8.90 |
| 9511 | 57.10 | 1.40 | 12.60 | 6.50 | 28.30 | 1.39 | 9.50 | 7.50 |
| 9513 | 79.30 | 1.08 | 73.70 | 18.80 | 47.80 | 1.06 | 12.00 | 9.00 |
| 9514 | 73.10 | 1.46 | 39.20 | 10.50 | 51.20 | 1.48 | 6.60 | 7.70 |
| 9515 | 80.00 | 1.29 | 43.00 | 11.20 | 54.60 | 1.40 | 8.50 | 7.80 |
| 9517 | 89.40 | 0.83 | 80.30 | 21.90 | 58.70 | 1.28 | 11.00 | 7.30 |
| 9518 | 73.60 | 1.40 | 32.00 | 8.80 | 48.30 | 1.46 | 8.20 | 7.40 |
| 9519 | 75.60 | 1.37 | 34.90 | 8.40 | 53.90 | 1.35 | 6.00 | 8.40 |
| 9520 | 66.60 | 1.49 | 24.00 | 7.80 | 47.90 | 1.41 | 6.00 | 8.90 |
| 9524 | 70.90 | 1.07 | 70.30 | 15.00 | 40.90 | 1.10 | 13.60 | 8.20 |
| 9525 | 56.70 | 1.19 | 66.60 | 14.00 | 31.10 | 1.21 | 15.40 | 8.40 |
| 9528 | 54.00 | 1.46 | 29.90 | 8.80 | 46.40 | 1.39 | 5.20 | 8.40 |
| 9529 | 47.80 | 1.45 | 24.10 | 9.30 | 40.30 | 1.33 | 6.20 | 9.10 |
| 9535 | 74.50 | 1.66 | 14.60 | 6.10 | 51.00 | 1.70 | 11.70 | 6.40 |
| 9565 | 96.20 | 1.05 | 93.10 | 27.40 | 68.60 | 1.20 | 10.20 | 8.60 |
| 9566 | 82.10 | 1.39 | 34.40 | 7.40 | 25.00 | 1.30 | 6.50 | 9.60 |
| 9567 | 67.80 | 1.17 | 23.70 | 8.10 | 61.70 | 1.47 | 12.00 | 8.00 |
| 9569 | 87.20 | 0.88 | 37.30 | 10.30 | 25.50 | 1.14 | 15.80 | 7.00 |
| 9570 | 90.60 | 1.08 | 89.00 | 21.80 | 67.00 | 0.97 | 14.90 | 10.50 |
| 9571 | 86.70 | 1.25 | 35.90 | 13.60 | 67.80 | 1.29 | 7.80 | 9.00 |
| 9572 | 86.80 | 0.99 | 67.80 | 12.60 | 36.40 | 1.07 | 8.00 | 7.70 |
| 9575 | 90.40 | 1.16 | 59.70 | 11.50 | 36.70 | 1.52 | 6.10 | 6.00 |
| Avg. | 76.93 | 1.18 | 49.83 | 14.05 | 45.73 | 1.29 | 12.29 | 8.16 |
